# Dynamic remodeling of the pancreas immune landscape in obesity

**DOI:** 10.1101/2025.07.14.664803

**Authors:** Alexey Koshkin, Kranthi Kiran Kishore Tanagala, Anna Eichinger, Michael Chait, Aoife Young, Shanila Shakil, Junichi Yoshikawa, Yosuke Sakamoto, Steven B. Wells, Xiaojuan Chen, Boris Reizis, Donna L. Farber, Stuart P. Weisberg

## Abstract

Obesity is a known risk factor for diseases of the pancreas, including diabetes, pancreatic cancer and pancreatitis, but mechanisms remain unclear. To elucidate how obesity impacts pancreatic immune homeostasis, we performed spatial, transcriptomic and functional profiling of human pancreatic immune cells from obese and non-obese organ donors. Obesity was associated with higher density of tissue resident memory T-cells (TRM) in the exocrine pancreas which display high cytotoxic functions and aggregated around macrophages. Single cell sequencing of pancreatic macrophages revealed two main subsets - FOLR2^+^ CD11c^−^ fetal-derived macrophages with pro-repair and immunoregulatory function and a FOLR2^−^ CD11c^+^ monocyte-derived macrophages with greater T-cell interactions and pro-inflammatory function. In obesity, the pancreatic macrophage landscape shifts to lower predominance of FOLR2^+^ CD11c^−^ macrophages and higher FOLR2^−^ CD11c^+^ macrophages which interact selectively with the TRM and inflamed exocrine epithelium. Together, these results identify macrophage-T cell circuits and immune epithelial interactions that fuel chronic pancreatic inflammation in obesity – a potential unifying mechanism for obesity-related pancreatic diseases.

## INTRODUCTION

Obesity affects >40% of adults in the United States and drives chronic inflammation, a root cause of many diseases. The pancreas, which plays a central role in digestion, metabolic sensing and control, is highly sensitive to obesity. Inflammation in the exocrine pancreas, increases the risk for pancreatitis and pancreatic cancer – debilitating diseases with few treatment options^1–7^. In obese mouse models, cytokines derived from macrophages can cause dysfunction of insulin-secreting endocrine islets, thereby contributing to the pathogenesis of type 2 diabetes^8^. Relative to the large burden of obesity-related pancreatic diseases, very little is known concerning the mechanisms by which obesity influences pancreatic immune regulation.

Macrophages are the predominant immune cell lineage in most tissues and play key roles in controlling tissue homeostasis and inflammation. Tissue macrophages include lineages derived from fetal and adult hematopoiesis, and mouse models have shown that the exocrine pancreas contains an equal mixture of fetal- and adult monocyte-derived macrophages^9,10^. Mouse studies in diverse organs, including heart, brain and pancreas indicate that fetal-derived macrophages have homeostatic and repair functions, whereas monocyte-derived macrophages are implicated in pathologic tissue remodeling, and inflammation^11–16^. In obesity, specialized lipid associated macrophages in adipose tissue, atherosclerotic lesions and the liver help to process excess lipids, yet they also contribute to chronic inflammation^17–20^. The macrophage populations in human pancreas and their responses to obesity are not well defined.

Most tissues also contain tissue resident memory T-cells (TRM) - derived from the effector memory T-cells (TEM) of primary immune responses - which have specialized tissue adaptations and site-specific functions^21,22^. In gastrointestinal organs, TRM are long-lived guardians against secondary infections, provide immunosurveillance against tumors and interface with innate immune cells to boost tissue immune responses ^23,24^. In addition to providing localized immune protection, TRM balance their cytotoxic and effector functions with immunoregulatory functions to avoid excessive inflammation and collateral tissue damage^25^. Indeed, failure of TRM regulation is implicated in many gastrointestinal inflammatory diseases^26,27^.

We previously showed that human pancreatic TRM have robust effector potential and unique phenotypic, and transcriptomic profiles using an organ donor tissue resource that has enabled numerous groundbreaking studies of human tissue immunity during homeostasis^22,25,27,28^. Close in situ interactions with pancreatic macrophages are a key feature of pancreatic TRM, and pancreatic macrophages provide a balance of checkpoint and co-stimulatory signals that maintain TRM in a poised but restrained state^27^. The impact of obesity on the landscape of pancreatic TRM and their regulatory interface with pancreatic macrophages has not been studied.

Our access to organ donor tissues enables studies that integrate demographic and anthropomorphic parameters with tissue immune profiling. We analyzed how increasing body mass index (BMI) impacts the landscape and regulation of pancreatic T-cells and macrophages across 68 adult organ donors integrating transcriptomic, spatial and functional analysis. Our results show that higher BMI is associated with elevated TRM density and cytotoxicity in the exocrine pancreas, and greater TRM-macrophage interactions. Single cell sequencing of pancreatic macrophages revealed two prominent subsets including FOLR2^+^CD11c^−^ fetal-derived immunoregulatory macrophages and FOLR2^−^CD11c^+^ monocyte derived pro-inflammatory macrophages. The FOLR2^−^CD11c^+^ macrophages show increased predominance with higher BMI and selectively interact with TRM and inflamed pancreatic epithelium. These findings define key aspects of pancreatic immune dysregulation in obesity that may drive chronic inflammation and pancreatic diseases.

## RESULTS

### Increased density of TRM and macrophages in human pancreas during obesity

Pancreatic disease risk increases with elevations in BMI – a key indicator of adiposity that is used to diagnose obesity ^29–31^. To define how obesity impacts the immune landscape of the exocrine and endocrine pancreas, we performed multiplexed imaging on pancreas tissue sections from adult organ donors (range 18-71 years, median age, 52 years) without underlying pancreatic disease (n=32) (Figure S1A). The BMI range in the cohort was 21-47 kg/m^2^, median 33 kg/m^2^, and 62.5% of donors were classified as obese (BMI ≥ 30 kg/m^2^)^32^.

Consistent with our previous results, macrophages were the highest density immune lineage across all compartments^27^ (Figure 1A, 1B). In the non-obese donors, macrophage and T-cell densities were highest in acinar areas, as compared to ductal, and lowest in islets (Figure 1B-1D). Macrophage densities were unchanged by obesity in acinar and ductal areas but were increased in islets (Figure 1B). In contrast, CD8 (Figure 1C) and CD4 (Figure 1D) T-cell densities were increased across all compartments in obesity, with the most marked increases in ductal CD8 T-cells (Figure 1C).

**Figure 1.**
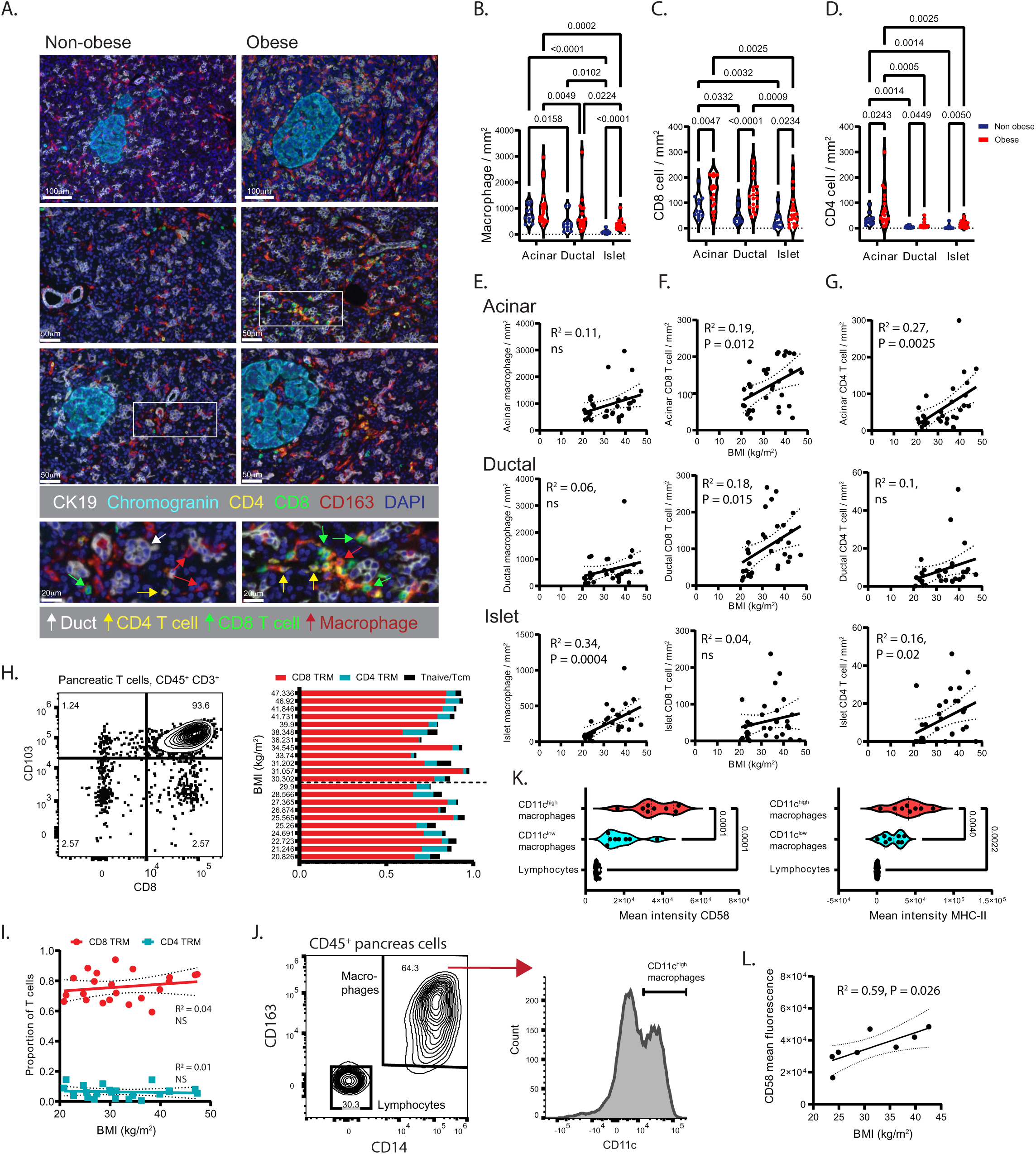
Quantification and phenotyping of the T-cells and macrophages in human pancreas and their associations with BMI. (A) Representative multiplex-stained sections of non-obese (left) and obese (right) pancreas are shown with markers for endocrine islets (chromogranin), ductal cells (CK19) and the immune lineages macrophages (CD163), and T cells (CD4, CD8). White boxes show areas with macrophages and T cells in the exocrine pancreas. The densities of (B) macrophages, (C) CD8 T cells and (D) CD4 T cells in the acinar, ductal and islet compartments are shown for obese (n=20, red circles) and non-obese (n=12, blue circles) organ donors. P value as calculated by two-way ANOVA with Tukey’s multiple comparisons test. The densities of (E) macrophages, (F) CD8 T cells and (G) CD4 T cells are plotted against donor body mass index (BMI) in the acinar (top), ductal (middle), and islet (bottom) compartments of pancreas (n=32 donors). (H) A representative contour plot is shown depicting CD8 and CD103 marker expression on pancreatic T-cells (CD45^+^ CD3^+^) (left). Shown also (right) are the donor BMI values with the quantification of CD8 (red) and CD4 (cyan) pancreatic tissue resident memory (TRM, CD45RA^−^ CCR7^−^ CD69 and/or CD103^+^) and the combined naïve (CD45RA^+^ CCR7^+^) and central memory (Tcm, CD45RA^−^ CCR7^+^) T-cell subsets as a proportion of total T-cells. (I) The proportions of pancreatic CD8 (red) and CD4 (cyan) TRM are shown plotted against donor BMI. (J) Shown is the flow cytometry gating strategy to identify human pancreatic macrophage (CD45^+^ CD14^+^ CD163^+^) (left) subsets with histogram showing CD11c expression (right) and gating used to identify CD11c^high^ macrophages. (K) The quantification is shown of CD58 (left) and MHC-II (right) mean fluorescence signal intensity for the indicated macrophage subsets and pancreatic lymphocytes in organ donors (n=8). (L) CD58 mean fluorescence is plotted against donor BMI (n=8). The best fit lines, 95% confidence intervals and P values were calculated using simple linear regression (ns, not significant).

We examined how immune lineages change as a function of BMI in each pancreatic compartment. By simple linear regression, BMI significantly correlated with higher macrophage density in islets (Figure 1E). BMI significantly correlated with higher CD8 T-cell density in acinar and ductal areas (Figure 1F); and higher CD4 density in acinar areas and islets (Figure 1G). These relationships were also found to be statistically significant by multivariable analysis after adjustment for known pancreas disease risk factors such as age, diabetes and being male (Table S1, S2). Thus, obesity directly correlates with distinct immune cell composition in exocrine and endocrine pancreas, with macrophage accumulation being most prominent in islets and T-cell accumulation most prominent in exocrine pancreas.

We assessed BMI-related changes in T-cell subset distribution by flow cytometry. Markers of tissue residency (e.g., CD69, CD103) on CD45RA^−^ CCR7^−^ TEM cells delineate TRM that are retained in, and specially adapted to, the tissue microenvironment ^27,33^. To this end, pancreatic T-cell subset distributions were analyzed in 22 adult donors with BMI of 21-47 kg/m^2^. CD8^+^ CD103^+^ TRM were the predominant pancreatic T-cell subset in obese and non-obese donors (Figure 1H) and TRM subset proportions did not correlate with BMI (Figure 1I).

Moreover, no BMI-related changes were observed in the minor populations of naïve (CD45RA^+^ CCR7^+^) and central memory (CD45RA^−^ CCR7^+^) T-cells (Figure 1I). Thus, the increased T-cell density in pancreas tissue found in obesity did not alter T-cell subset distributions, with CD8 TRM as the predominant pancreatic T cell subtype.

Pancreatic macrophages, identified as CD14^+^CD163^+^ from our previous work^27^, showed heterogeneity in expression of the pro-inflammatory marker CD11c (Figure 1J). The CD11c^high^ macrophages expressed the highest surface levels of CD58, a T-cell adhesion and co-stimulation molecule ^34^ and the antigen presentation molecule MHC-II compared to CD11c^low^ macrophages (Figure 1K). Moreover, CD58 expression on CD11c^high^ macrophages positively correlated with donor BMI (Figure 1L). Thus, CD11c^high^ pancreatic macrophages express molecules that promote T-cell adhesion and activation, including CD58 which is elevated in obesity.

### T-cells aggregate around CD11c^high^ macrophages in the exocrine pancreas of obese donors

We used multiplex imaging to analyze the spatial relationships of macrophages and T cells in the exocrine and endocrine pancreas. Macrophages were classified as either CD11c^high^ or CD11c^low^ (Figure 2A, Figure S2A) and the CD11c^high^ macrophages were found to be markedly enriched around the CK19^+^ ductal epithelium relative to acinar or islet areas (Figure 2B, S2B). The lowest proportions of CD11c^high^ macrophages were in islets with intermediate levels in the acinar areas (Figure 2B). Moreover, donor BMI significantly correlated with the proportion of CD11c^high^ macrophages around the CK19^+^ ductal epithelium but not within acinar or islet areas (Figure 2C, 2D, 2E). Most of the pancreatic T-cells clustered within 20μm of the CD11c^low^ macrophages, however larger clusters of T-cells formed around CD11c^high^ macrophages in obese donors (Figure 2F, 2G) and the percentage of CD8 T-cells located within 20μm of the CD11c^high^ macrophages positively correlated with donor BMI (Figure 2H).

**Figure 2.**
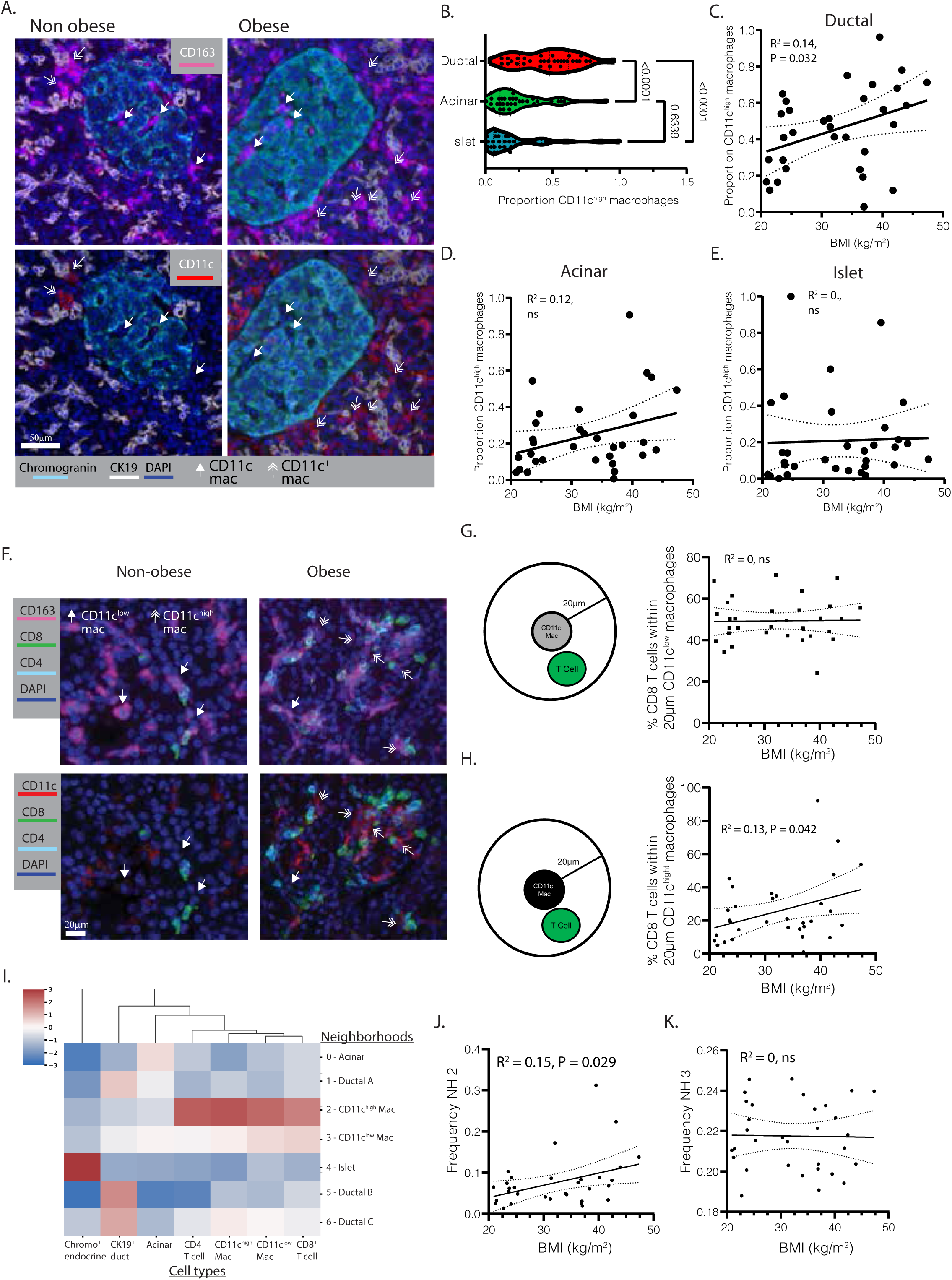
Altered macrophage T-cell interactions in human pancreas with increasing BMI. (A) Representative multiplex fluorescence imaging of human pancreatic tissue sections, depicting CD163 staining (top, magenta) and CD11c staining (bottom, red) in paired images with pancreatic ducts labeled by CK19 (white), and islets indicated by chromogranin (cyan). Solid arrowheads indicate CD11c^low^ macrophages whereas double-line arrowheads indicate CD11c^high^ macrophages. (B) Violin plots quantifying the CD11c^high^ macrophages as a proportion of total macrophages within ductal, acinar and islet pancreas tissue compartments. P values were calculated by one-way ANOVA and Holm-Sidak multiple comparison test. (C-E), Scatter plots show correlations between the proportions of CD11c^high^ macrophages within (C) ductal, (D) acinar, and (E) islet niches (y-axis) and donor BMI (x-axis; n = 32). (F) Multiplex fluorescence images showing spatial interactions among T-cells and macrophages depicting CD163 staining (top, magenta) and CD11c staining (bottom, red) in paired images with CD8 (green) and CD4 (cyan) T-cells. (G, H) Scatter plots show the relationship between donor BMI (x-axis, n = 32) and the percentage of CD8 T cells within a 20µm radius of either (G) CD11c^low^ or (H) CD11c^high^ macrophages. (I) Heatmap of seven pancreatic cellular neighborhoods (NHs), characterized by differential enrichment of the seven indicated cell types (CTs, bottom). NH classification (right) was derived from pooled data across all donors (n = 32) and named according to lineage enrichment. Color intensity reflects the relative enrichment score of each cell type within each NH. (J, K) Scatter plots show the correlation between BMI and frequency of (J) NH2 and (K) NH3. The best fit lines, 95% confidence intervals and P values were calculated using simple linear regression.

We further analyzed the spatial orientation of T-cells and macrophages in pancreas by defining 7 ‘neighborhoods’ (NH) each with distinct enrichment of spatially associated cell lineages (Figure S2C). Non-immune cell NH included those with high density of chromogranin^+^ endocrine cells (NH4) and CK19^+^ ductal cells (NH1, 5, 6) (Figure 2I)^35^. Pancreatic macrophages were enriched in NH2 and 3 (Figure 2I). The CD11c^high^ macrophages, along with CD4 and CD8 T-cells and CD11c^low^ macrophages, were most strongly enriched in NH2 (Figure 2I) and NH2 frequency uniquely correlated with donor BMI (Figure 2J, 2K). Thus, T-cells accumulate as discrete foci clustering around CD11c^high^ macrophages in the exocrine pancreas of obese donors.

### Pancreatic TRM express transcriptome signatures of high cytolytic effector function

We analyzed tissue specific transcriptomic signatures of pancreatic TRM by performing Cellular Indexing of Transcriptomes and Epitopes by Sequencing (CITE-seq) on T-cells purified from the pancreas, pancreas draining lymph nodes (PLN), small intestine, and spleen of adult organ donors. Cluster analysis showed substantial T-cell transcriptomic variation associated with differentiation state and anatomic site (Figure 3A, Figure S3A). Clusters with low effector molecule expression (e.g., *TBX21*, *GZMA*, *CCL5*) (Figure S3B) showed high expression of *CCR7,* the lymphoid tissue homing receptor (Figure 3B) and transcription factor *TCF7* (Figure S3B), consistent with naïve and central memory T-cells (clusters #4, 9, 10); and these were predominately derived from lymph node and spleen (Figure S3A). Most cells in the remaining clusters expressed CD8 and high levels of CD103, consistent with TRM (Figure 3B). These TRM were separated into two distinct tissue specific groups: those derived from pancreas (clusters 0, 2, 5) and those derived from small intestine (clusters 1, 3, 6, 7, 11; Figures 3A and S3A). T-cells derived from islet fractions distributed into pancreas-enriched clusters (clusters 0, 2, 5) and clusters 8 and 9 containing effector T-cells from PLN, spleen and pancreas (Figure S3A).

**Figure 3.**
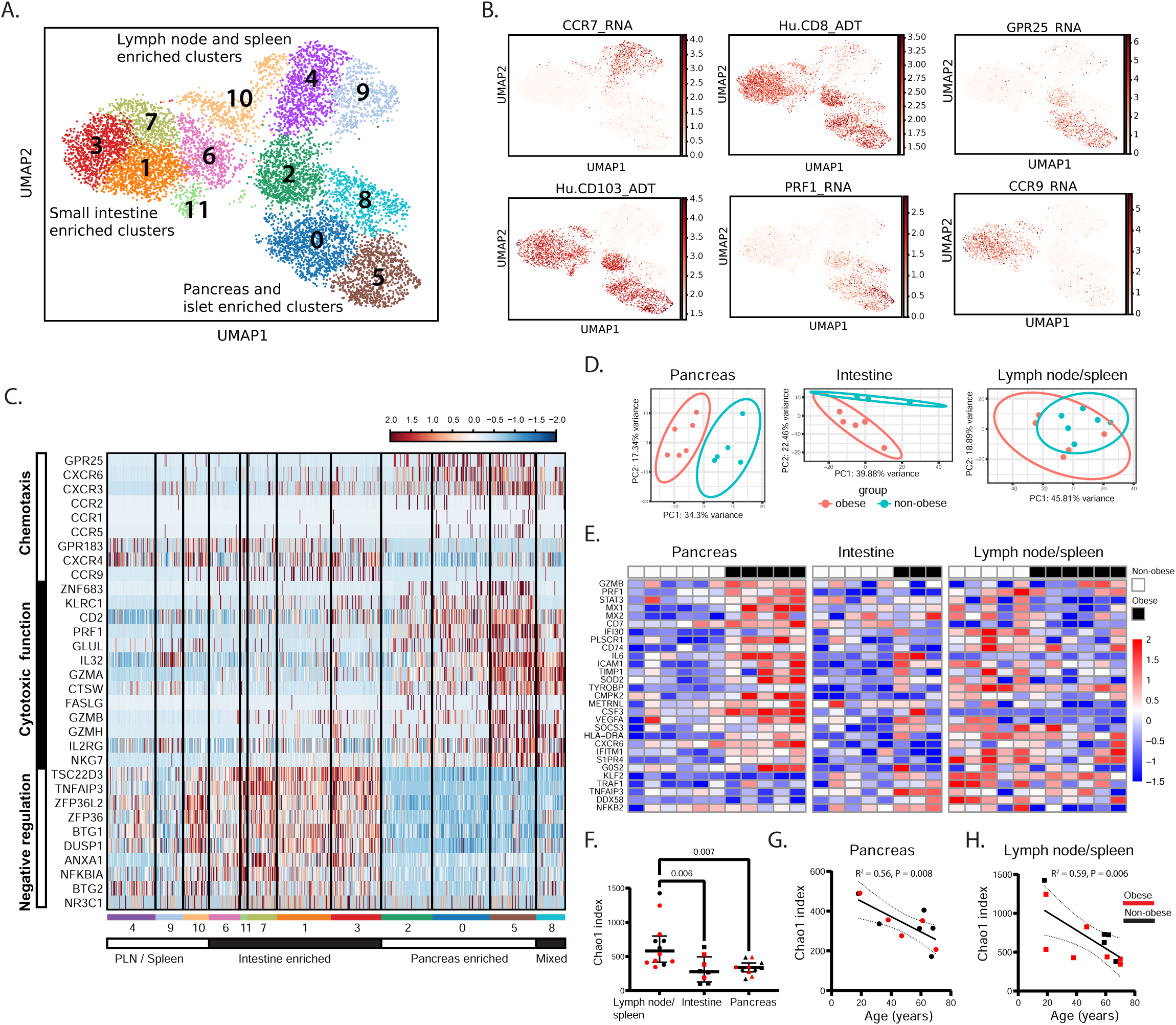
Pancreatic tissue resident memory T cells express tissue-specific gene signatures of high effector function and show unique gene expression changes in obesity. (A) Shown are the Uniform Manifold Approximation and Projection (UMAP) embeddings and cluster analysis of total T cells obtained from pancreas, purified islets, pancreas draining lymph node (lymph node), intestine and spleen of 1 non-obese and 2 obese organ donors with numbers and colors indicating distinct clusters. The observed tissue enrichment of the clusters is indicated. (B) Normalized values for the indicated RNAs and antibody-derived tags (ADT) are depicted on the scaled feature plots. (C) Differentially expressed genes (DEG) in the pancreas-enriched T-cell clusters (0, 2, 5) are shown in the normalized and scaled heatmap with color-coded bars below corresponding to clusters identified in the UMAP and organized according to tissue enrichment. The functional categories of the genes based on ontology analysis are indicated (left). (D) Principal component analyses with 95% confidences ellipses are shown for bulk RNA sequencing data of sorted CD8 TRM from pancreas (left), intestine (middle) and lymph node/spleen (right) from obese (n=5) and non-obese (n=6) organ donors. (E) DEG in the ‘interferon’ and ‘inflammatory response’ and ‘allograft rejection’ gene sets that are significantly increased in pancreas TRM from obese versus those from non-obese donors (P adj < 0.05) are shown on the normalized and scaled heatmap. (F) Shown is the Chao1 T-cell receptor (TCR) diversity metric of the CD8 TRM from combined lymph node/spleen (n=11), intestine (n=8) and pancreas (n=11) for obese (red symbols) and non-obese (black symbols) donors. P values were calculated using the Kruskal-Wallis and Dunn’s multiple comparisons test. (G) The Chao1 index for the CD8 TRM from pancreas (n=11) and (H) lymph node/spleen (n=11) is plotted against donor age with red symbols indicating obese donors. The best fit lines, 95% confidence intervals and P values were calculated using simple linear regression.

In pancreas-enriched TRM clusters, there were 523 differentially expressed genes (DEG). Transcripts with higher expression in the pancreas TRM were enriched in pathways related to chemokine signaling, and cytotoxic T cell function. Transcripts with lower expression in pancreatic TRM were enriched in pathways related to immunoregulation (Table S3)^36^. Unique chemokine receptor profiles also distinguished the pancreatic TRM including high expression of the CXCL17 receptor, *GPR25*^37^, and the CXCL16 receptor, *CXCR6*^38^, and low expression of the *CCR9* intestinal T-cell homing receptor^39^ (Figure 3B, 3C). Pancreatic TRM also showed the highest expression of transcripts implicated in cytotoxic function, including the ZNF683 transcription factor^40^, multiple effector molecules (i.e., *PRF1*, *GZMA*, *GZMB*, *FASLG*) and the pro-inflammatory, pro-tumorigenic *IL32* cytokine^41^ (Figure 3B, 3C). In addition, the transcript encoding CD2, the T-cell adhesion and co-stimulatory receptor for CD58, showed the highest expression in pancreatic TRM (Figure 3C). Conversely, multiple transcripts involved in negative regulation of T-cell function were markedly decreased in pancreatic TRM, including *TNFAIP3* and *NFKBIA*, which encode the NF-κB signaling inhibitors A20 and IκBα, *NR3C1*, encoding the glucocorticoid receptor, and glucocorticoid-induced genes (e.g. *TSC22D3*, *ZFP36*, *BTG1*) (Figure 3C, Figure S3C). By contrast, the intestinal TRM showed markedly lower levels of cytotoxic and effector genes and higher expression of immunoregulation genes (Figure 3C). Thus, pancreatic TRM express a unique transcriptome signature of pancreas-specific trafficking, macrophage adhesion, and high cytotoxic effector function.

### Pancreatic TRM show tissue-specific pro-inflammatory changes in obesity

Obesity-related gene expression changes were analyzed by bulk RNA sequencing in sorted CD8^+^ CD69^+^ TRM from pancreas, intestine, PLN, and spleen from 6 non-obese (BMI 22.7 – 26.8 kg/m^2^) and 5 obese (BMI 34.5 – 46.9 kg/m^2^) organ donors (Figure S3D), including 3 donors from a prior study (GSE135582)^27^. As expected, the TRM transcriptomes varied by tissue site (not shown). Principal component analysis (PCA) within tissue sites showed that pancreatic and intestinal TRM cluster by obesity status while TRM from the lymphoid tissues (PLN and spleen) do not cluster by obesity status (Figure 3D). Differential gene expression analysis showed higher numbers of obesity-related DEGs in the pancreatic TRM (738 genes) compared to those from the other tissue sites (lymph node and spleen, 5 genes; intestine 402 genes). The pancreatic TRM DEGs that were increased in obesity showed the greatest enrichment in pathways related to interferon response (e.g. *STAT3*, *MX1*, *MX2*, *IFI30*), inflammation (e.g. *CSF3*, *ICAM1*, *CXCR6*) and T-cell effector function (e.g. *PRF1*, *GZMB*) (Figure 3E, Table S4)^36^. In contrast, intestinal TRM DEGs that were increased in obesity have less enrichment in these pathways and some function as brakes on inflammation (e.g. *TNFAIP3*, *DDX58*^42^) (Figure 3E). Thus, pancreatic TRM uniquely exhibit altered gene expression in obesity consistent with a pro-inflammatory state.

To determine if the increased density and gene expression changes of pancreas TRM in obesity are related to clonal expansion, TCR sequences were extracted from the RNA sequencing datasets^43^. Plotting the clonal diversity metric (i.e., the Chao1 index) against tissue site showed lower diversity in TRM from pancreas and small intestine compared to TRM from lymphoid tissues (i.e., PLN, spleen); TRM diversity in each tissue was unchanged in obese compared to non-obese, donors. (Figure 3F). In addition, plotting clonal diversity against age showed significant aging-related decline in clonal diversity for pancreatic (Figure 3G) and lymphoid (Figure 3H) TRM without any effect of BMI. Thus, the increased density of pancreatic TRM in obesity is not linked to clonal expansion.

### A differentiation trajectory towards CD11c^high^ macrophages from infiltrating monocytes in human pancreas

To identify transcriptomic signatures underlying phenotypically distinct pancreatic macrophages, we performed CITE-seq from 5 organ donors without pancreatic disease (3 non-obese, 2 obese). Cluster analysis identified three main macrophage populations (Figure 4A). 1) Cluster 0 showed extensive similarity with fetal -derived pancreatic macrophages in murine fate mapping studies^15,16^, including *FOLR2*, *CD209*, *MRC1* (CD206), and *LYVE1* (Figure 4B, 4C, S4A, S4B). These macrophages also express the highest levels of the efferocytosis molecules *MERK*, *AXL*, and *GAS6*, which play key roles in immunoregulation during tissue repair^44^ (Figure 4B, S4B). 2) Monocytic macrophages (clusters 3, 5, 6) selectively express the monocyte chemotaxis receptor, CCR2 (Figure 4B, 4C, S4B), along with genes associated with bone marrow derived monocytes (e.g. *S100A12*, *FCN1, LYZ*^45^), monocyte migration into tissues (*CCR1, SELL*, *CD44*), and the CD36 phagocytic receptor and fatty acid transporter (Figure 4B, S4B). 3) CD11c^high^ macrophages (clusters 1, 2, 4) have low expression of fetal-derived macrophage markers and monocytic markers (Figure 4B, 4C, S4B) and have high expression of the lipid-associated macrophage markers *SPP1*, *TREM2*, and *CD9* (Figure 4B, 4C, S4A, S4B). Cluster 2 of CD11c^high^ macrophages is particularly enriched in genes associated with lysosome function and lipid metabolism (Figure 4B, S4B). Clusters 1 and 4 of CD11c^high^ macrophages expressed the highest levels of CD58, and the pro-inflammatory marker, Podoplanin (PDPN)^46–51^ (Figure 4B, S4B). Clusters of cycling macrophages contained both FOLR2^+^ (cluster 10) and CD11c^+^ (clusters 7, 8, 9) subsets (Figure 4A-4C). Thus, CD11c expression delineates a pro-inflammatory macrophage subset that is transcriptionally and phenotypically distinct from the FOLR2^+^ macrophage subset.

**Figure 4.**
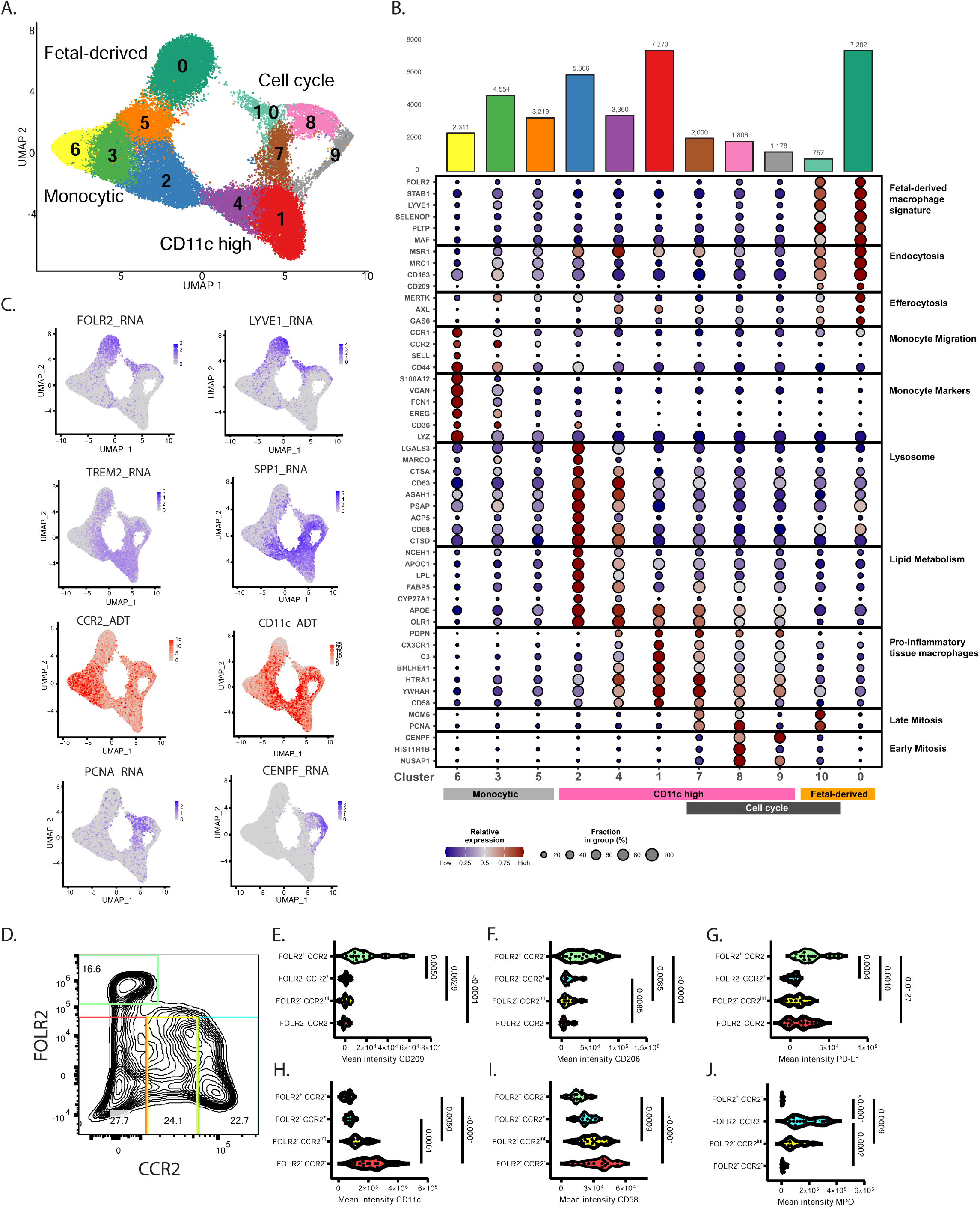
Integrated transcriptomic and phenotypic profiling of pancreatic macrophages from obese and non-obese organ donors. (A) Uniform Manifold Approximation and Projection (UMAP) visualization of pancreatic macrophage populations derived from single cell analysis of samples from non-obese (n = 3) and obese (n = 2) donors. Clusters are indicated by colors and numbers. Classification of the macrophage clusters based on gene expression and phenotype is indicated. (B) Shown is a dot plot in which dot size corresponds to the fraction of cells in each cluster expressing the indicated gene and color corresponds to relative RNA expression scaled by row. The cluster color from the UMAP and cell number are indicated along the top. Cluster numbers are indicated along the bottom with macrophage classifications based on phenotype and gene expression, and the genes are organized by function (right). (C) Feature plots show the normalized expression of the indicated cluster marker genes (purple), and surface proteins (red) with expression level corresponding to color intensity for individual cells on the UMAP. (D) Representative flow cytometry contour plot of pancreatic macrophages showing FOLR2 and CCR2 expression. Classification gates are indicated by the colored boxes including FOLR2^+^CCR2^−^ (green), FOLR2^−^CCR2^+^ (cyan), FOLR2^−^CCR2^int^ (yellow) and FOLR2^−^CCR2^−^ (red). (E-J) Compiled data from 13 non-obese and obese organ donors showing mean intensities (x-axis) of the indicated markers across the indicated macrophage subsets (y-axis) for (E) CD209, (F) CD206, (G) PD-L1, (H) CD11c, (I) CD58 and (J) MPO. P values were calculated by one-way ANOVA and Holm-Sidak multiple comparison test.

Pseudo-temporal trajectory inference was used to assess the lineage relationships of the FOLR2^−^ macrophage subset (clusters #1-9) (Figure S4C)^52^ . Monocyte markers and transcripts involved in monocyte migration from blood into tissue (e.g. *S100A9*, *CCR1*, *CCR2*) were highly expressed in cells at the beginning of the trajectory and declined with trajectory progression (Figure S4C). In contrast, CD11c surface expression were increased over trajectory progression along with markers of lipid-associated macrophages (e.g. *TREM2*, *CD9*, *SPP1*), genes involved in T-cell activation (e.g. *HLA-DRA*, *CD86*, *CCL4, IL18*) and cell cycle genes (Figure S4C). These results provide a model for how infiltrating monocytic macrophages may undergo functional specialization in the pancreas where increased surface expression of CD11c is linked to acquisition of lipid-associated macrophage markers, entry into the cell cycle, and increased capacity for T-cell activation.

We validated the subsets identified by CITE-seq using flow cytometry. Pancreatic macrophages (CD45^+^ CD14^+^ CD163^+^ cells) from 5 non-obese and 8 obese organ donors were separated based on expression of FOLR2 and CCR2 (Figure 4D). FOLR2^+^ CCR2^−^ (Figure 4D) macrophages expressed high levels of the CD206 and CD209 scavenger receptors and the checkpoint ligand PD-L1 (Figure 4E-4G) and had the lowest levels of CD11c (Figure 4H) and CD58 (Figure 4I). The FOLR2^−^ macrophages separated into 3 subsets expressing high, intermediate, and low levels of CCR2 (Figure 4D). The FOLR2^−^ CCR2^high^ macrophages contained the highest levels of intracellular myeloperoxidase (MPO) (Figure 4J) - a specific marker of bone marrow myeloid progenitors and monocytes that declines with myeloid differentiation. Within the FOLR2^−^ macrophage lineage, decreased CCR2 expression was associated with lower intracellular MPO (Figure 4J), and increased CD11c (Figure 4H) and CD58 (Figure 4I). Thus, pancreatic macrophages functionally segregate based on their ontogeny and differentiation in pancreas, with FOLR2^+^ CD11c^−^ phenotype indicating probable fetal-derived macrophages with immunoregulatory functions and FOLR2^−^ CD11c^+^ phenotype delineating monocyte-derived pro-inflammatory macrophages.

### Altered pancreatic macrophage subset composition in obesity enhances TRM functions

We analyzed how BMI associates with pancreatic macrophage subset distributions across two main phenotypes: FOLR2^+^ CD11c^−^ and FOLR2^−^ CD11c^+^ (Figure 5A). Subset quantification as a proportion of the total (CD14^+^ CD163^+^) macrophage population showed that increasing BMI is correlated with decreased FOLR2^+^ CD11c^−^ macrophages (Figure 5B) and increased FOLR2^−^ CD11c^+^ macrophages (Figure 5C). Thus, the pancreatic macrophage landscape shifts in obesity, with pro-inflammatory CD11c^+^ macrophages displacing fetal-derived immunoregulatory FOLR2^+^ macrophages.

**Figure 5.**
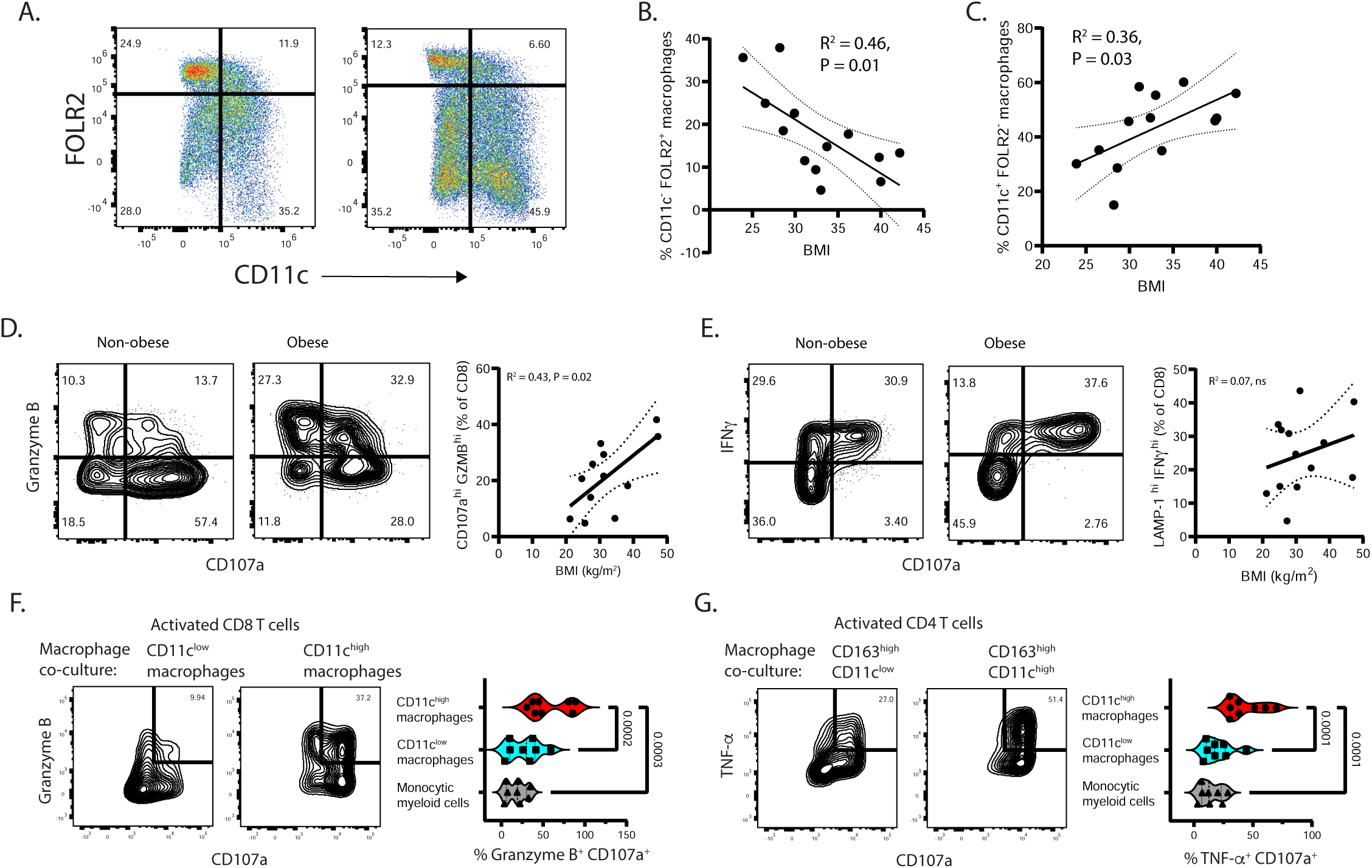
Alterations of pancreatic macrophage subsets with increasing BMI and their effects on TRM function. (A) Representative pseudo color plots of pancreatic macrophages from a non-obese (left) and obese (right) organ donor showing gates used to classify FOLR2^+^CD11c^-^ and FOLR2^-^CD11c^+^ macrophage populations. (B-C) Scatterplots are shown correlating the percentages of FOLR2^+^CD11c^-^ (B) and FOLR2^-^CD11c^+^ (C) macrophage subsets versus donor BMI. (D, E) Contour plots of pancreatic CD8 TRM after overnight autologous pancreatic macrophage co-culture and 4-6 hour activation plotting intracellular Granzyme B (GZMB) (D) and interferon-γ (IFNγ) (E) versus the surface degranulation marker CD107a from a representative non-obese (left) and obese (middle) donor. Compiled data showing the percentages of GZMB^+^ CD107a^+^ cells versus donor BMI (n=12) (D) and IFNγ^+^ CD107a^+^ cells versus donor BMI (n=13) (E) are shown to the right of the contour plots. The best fit lines, 95% confidence intervals and P values were calculated using simple linear regression analysis. (F, G) Flow cytometry data from naïve CD8 (F) and CD4 (G) T-cells after activation and 72-hour co-culture with CD11c^low^ and CD11c^high^ macrophages. Representative contour plots (left) show intracellular Granzyme B versus surface CD107a of CD8 T-cells (F) and intracellular TNF-α versus surface CD107a of CD4 T-cells (G). The compiled data are shown to the right of the contour plots quantifying the percentages of Granzyme B^+^ CD107a^+^ CD8 T-cells (F) and the percentages of TNF-α^+^ CD107a^+^ CD4 T-cells (G) (n=8). P values were calculated by one-way ANOVA and Holm-Sidak multiple comparison test.

We hypothesized that the altered pancreatic macrophage landscape in obesity can influence the function of TRM, which our previous work showed is shaped by pancreatic macrophages^27^. We examined the relationship of BMI with the functional profiles of pancreatic TRM after activation in autologous pancreatic macrophage co-culture. In this setting, significant proportions of TRM showed concomitant CD107a degranulation and high expression of the Granzyme B cytotoxic molecule (CD107a^+^ GZMB^+^ cells, 21.5±3.5%) (Figure 5D) and interferon γ (CD107a^+^ IFN γ ^+^ cells, 24.5±3.2%) (Figure 5E). The proportions of CD107a^+^ GZMB^+^ cells, but not of the CD107a^+^ IFN γ ^+^ cells, significantly correlated with donor BMI by linear regression (Figure 5D, Figure 5E). This enhanced cytotoxic function of pancreatic TRM associated with BMI may reflect their increased interactions with CD11c^+^ pro-inflammatory macrophages.

We tested mechanisms of pancreatic macrophage-T-cell interaction using a co-culture assay in which naïve T-cells are activated by monomeric anti-CD3 antibody with sorted pancreatic macrophages (Figure S5A). Flow cytometry showed that pancreatic macrophages express co-stimulatory ligands, CD86 (Figure S5B) and CD58 (Figure 1K), that enhance T-cell effector functions along with TGF-β (Figure S5B) which promotes tissue residency molecule expression ^53–55^. After activation and co-culture with pancreatic macrophages, naïve CD8 T-cells showed strong induction of CD103 and GZMB (Figure S5C, S5D) and naïve CD4 T-cells showed TNF-α production (Figure S5E). By contrast, T-cell activation without macrophages (using anti-CD3/28/2 beads) did not induce CD103 (Figure S5C). Antibodies blocking TGF-β prevented T-cell expression of CD103 in macrophage co-culture but did not affect GZMB (Figure S5C, S5D). Antibodies blocking CD58 reduced both GZMB and CD103 induction (Figure S5C-S5D); and inhibition of CD86 with CTLA-4-Ig^55^ prevented GZMB and TNF-α production (Figure S5D, S5E). Thus, pancreatic macrophages drive T-cell residency and effector molecule expression by providing TGF-β with co-stimulatory ligands.

The pro-inflammatory profile of CD11c^+^ macrophages and high CD58 expression (Figure 1K) may boost their T-cell interactions, so we compared the capacities of sorted CD11c^+^ and CD11c^−^ pancreatic macrophage subsets to drive effector and residency molecule induction in naïve T-cells. Both subsets similarly promoted CD103 expression on naïve CD8 T-cells (Figure S5F) after activation and co-culture, but the CD11c^high^ macrophages drove higher CD8 T-cell GZMB production and CD107a degranulation (CD107a^+^ GZMB^+^) (Figure 5F) and significantly higher expression of cytotoxic perforin (Figure S3G). The CD11c^high^ macrophage subset also promoted higher CD4 T-cell TNF-α production and CD107a degranulation (CD107a^+^ TNF-α^+^) (Figure 5G). These data show that CD11c^high^ pancreatic macrophages have the greatest capacity to drive T-cell effector functions.

### Macrophages and T-cells in the pancreas of obese donors organize around inflamed pancreatic epithelium

To gain insight into the spatial arrangement of the pancreatic macrophage subsets and their interactions with pancreatic epithelium, multiplex staining of pancreas tissue microarrays from obese (n=14) and non-obese (n=14) donors was performed using a custom panel of epithelial and immune lineage markers (see Methods, Table S8). Consistent with the macrophage CITE-seq data (Figure 4B, 4C) and flow cytometry validation (Figure 4D-4J) a subset co-express MPO and CD36 (Figure S6A, S6B), consistent with the phenotype of monocytic macrophages (Figure 4B, S4B, 4J). Among MPO^−^ macrophages, FOLR2/CD209^+^ CD11c^−^ macrophages expressed the highest levels of CD163 (Figure S6A, S6C), an anti-inflammatory marker^56^ highly expressed by fetal-derived macrophages (Figure 4B, S4B); whereas FOLR2/CD209^−^ CD11c^+^ macrophages expressed the lowest levels of CD163 and highest levels of MHC-II (Figure S6C, S6D). Moreover, the FOLR2/CD209^−^ CD11c^+^ macrophages positively correlated, and FOLR2/CD209^+^ CD11c^−^ macrophages negatively correlated, with BMI (Figure 6A) independent of relevant covariates (Table S5). Nuclear Ki67 was higher in CD11c^+^ and FOLR2/CD209^+^ macrophages, as compared to the MPO^+^ macrophages (Figure S6E), consistent with their progression through the cell cycle as observed in the CITE-seq dataset (Figure 4A-4C, S4B).

**Figure 6.**
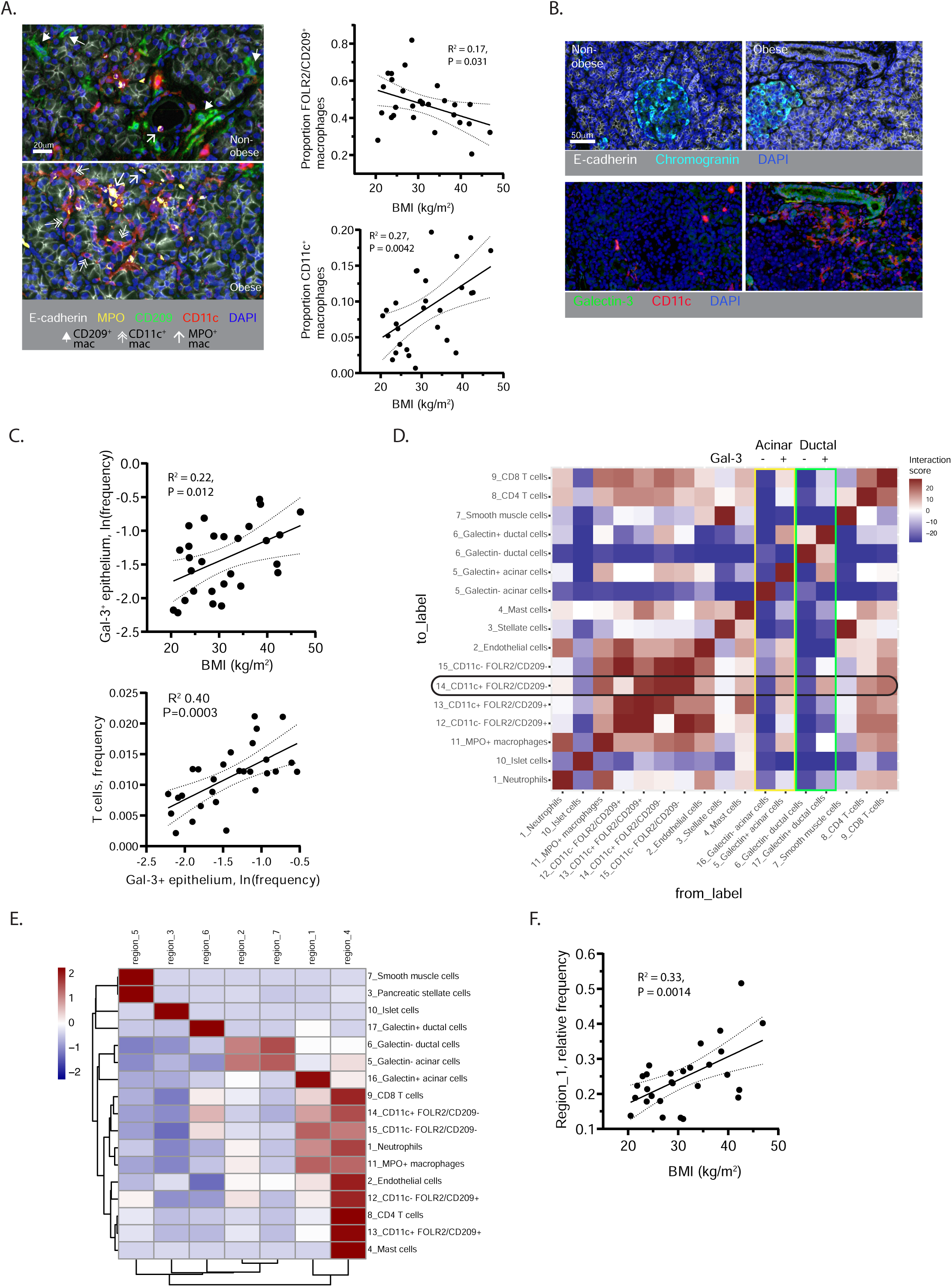
The spatial interactions between epithelial cells and immune microenvironment in human pancreas. (A) Representative images of multiplex stained pancreas from a non-obese (top) and an obese (bottom) organ donor showing epithelial cells (E-cadherin^+^, white), myeloperoxidase-positive monocytic macrophages (MPO^+^, single line arrow), CD11c^−^ FOLR2/CD209^+^ macrophages (solid arrow), and CD11c^+^ FOLR2/CD209^−^ macrophages (double line arrow). The scatterplots (right) show the proportions of CD11c^−^ FOLR2/CD209^+^ macrophages (top) and CD11c^+^ FOLR2/CD209^−^ macrophages (bottom) correlated with donor BMI (n=28). (B) Representative images of multiplex stained pancreas from a non-obese (left) and an obese (right) organ donor showing epithelial (E-cadherin) and islet (chromogranin) markers (top) with Galectin-3 (Gal-3) (green) and CD11c (red) (bottom). (C) Shown are scatterplots of Gal-3^+^ epithelial cells as a proportion of total pancreatic cells correlated with donor BMI (top) and of T cell frequency correlated with Gal-3^+^ epithelial cells frequency (bottom) (n=28). (D) A matrix heatmap (n=28 donors) shows cell-to-cell interaction scores reflecting the probability of spatial interaction where red indicates significantly increased interaction, and blue indicates significantly decreased interaction compared against random null distributions using permutation testing. A horizontal black oval marks CD11c^+^FOLR2/CD209^−^ macrophages. Yellow and green vertical boxes mark acinar (E-cadherin^+^cytokeratin^−^) and ductal (E-cadherin^+^cytokeratin^+^) cells respectively with Gal-3 expression indicated at the top. (E) Heatmap showing enrichment of 17 distinct cell types (right, rows) across 7 regions (top, columns) identified using the Local Indicators of Spatial Association analysis on multiplex stained pancreas tissue microarrays (n=28 donors). (F) The scatterplot shows frequencies of region 1 (y-axis) correlated with donor BMI (x-axis). The best fit lines, 95% confidence intervals and P values were calculated using simple linear regression analysis.

In exocrine pancreas of obese organ donors, E-cadherin^+^ Chromogranin^−^ epithelium showed higher expression of the pro-inflammatory adhesion molecule Galectin-3 (Gal-3), which is known to be upregulated in obesity, chronic pancreatitis and pancreatic cancer ^57–62^(Figure 6B). The proportion of Gal-3^+^ E-cadherin^+^ epithelial cells relative to total pancreas cells significantly correlated with donor BMI (Figure 6C); in a multivariable model, this relationship was independent of relevant co-variates (Table S6). Pancreatic T-cell density in this dataset also correlated with donor BMI (Table S6, Figure S6F), as in the previous dataset (Figure 1F, 1G), and with Gal-3^+^ epithelial cell density (Figure 6C), suggesting a link between pancreatic epithelial inflammation and T-cell accumulation.

To determine how Gal-3^+^ exocrine epithelial cells are spatially related to the pancreatic immune cells, the probability of pairwise interaction between cell types (at 20μm radius) was quantified^63^(Figure 6D). Macrophages, T-cells, and endothelial cells had high levels of mutual interaction in pancreas (Figure 6D). Notably, Gal-3^−^ ductal (E-cadherin^+^ CK19^+^ chromogranin^−^) and acinar (E-cadherin^+^ CK19^−^ chromogranin^−^) cells had little interaction with immune cells, whereas the Gal-3^+^ ductal and acinar cells had higher interactions with macrophages and T-cells, and the greatest interactions with CD11c^+^ macrophages (Figure 6D).

To better define the spatial organization of pancreatic immune and epithelial cells, Local Indicators of Spatial Association (LISA) was used to define 7 regions with distinct enrichment of 17 different cell types in close spatial proximity (10-50μm) (Figure 6E)^64^. All pancreas immune lineages including macrophages, T-cells, mast cells, and neutrophils were highly enriched with endothelial cells in region #4. Endocrine cells and Gal-3^−^ epithelial cells were enriched in regions lacking immune cells (#2, 3, 7). In contrast, Gal-3^+^ acinar and ductal cells were enriched in regions 1 and 6 with CD11c^+^ CD209/FOLR2^−^ macrophages. Notably, the frequency of region 1, which is most enriched for Gal-3^+^ acinar cells, CD11c^+^ CD209/FOLR2^−^ macrophages, MPO^+^ macrophages, neutrophils, and CD8 T-cells (Figure 6D), positively correlated with BMI (Figure 6F). Thus, pro-inflammatory immune interactions of macrophages and T cells are spatially organized around inflamed Gal-3^+^ epithelium.

## DISCUSSION

In this study we identified key cellular mechanisms by which obesity drives chronic inflammation in the pancreas, a central metabolic sensor and primary tissue site implicated in obesity-associated diseases. Integration of CITE-seq with functional studies and spatial analysis showed how the interactions between pancreatic macrophages and TRM are reshaped with increasing BMI to establish self-sustaining inflammatory circuits. The selective interaction of inflamed Gal-3^+^ epithelium with macrophages and T-cells highlights how obesity drives pro-inflammatory epithelial-immune microenvironments. These findings substantially expand our understanding of the pancreatic response to obesity^65,66^ and lay the groundwork for strategies to mitigate chronic pancreatic inflammation and prevent pancreatic disease.

We previously identified CD8 TRM and macrophages as the predominant immune cells in pancreas which cluster together in the exocrine regions^27^. Here we show how obesity significantly alters the pancreatic immune architecture via compartment-specific reorganization. In the exocrine pancreas, TRM accumulate with increasing BMI (Figure 1F-1I) and are spatially organized around CD11c^+^ macrophages (Figure 2F-2J). The selective association of TRM-macrophage foci with inflamed Gal-3^+^ epithelium shows how immune-epithelial crosstalk fuels exocrine inflammation in obesity (Figure 6D-6F). Pancreatic islets show increased macrophage density with obesity which mirrors findings in mouse models and may reflect macrophage functions in supporting homeostatic islet expansion to increase insulin output in obesity^8^.

Our CITE-seq results provide novel insights into the transcriptomic states underlying the functions and interactions of pancreatic TRM and macrophages. Pancreatic TRM adopt a uniquely cytotoxic posture (high *GZMB*, *PRF1*, and *IL32*)^27,67^ while simultaneously downregulating multiple NF-κB brakes (*TNFAIP3*, *NFKBIA*, and the *NR3C1*-encoded glucocorticoid receptor)^68–72^(Figure 3B, 3C). The readying of TRM cytotoxic machinery while lowering anti-inflammatory checkpoints may predispose TRM to inappropriate activation. We also identified mechanisms driving TRM localization to pancreas including upregulation of GPR25, receptor for CXCL17^37,73^, and CXCR6, receptor for CXCL16^74^– key chemo-affinity systems for lymphocyte homing and retention in non-intestinal mucosal tissues (Figure 3B, 3C). The concurrent low expression of the intestinal-homing receptor, CCR9, by these pancreatic TRM^27^, indicates a dedicated pancreatic and non-intestinal mucosal-like tissue residency program (Figure 3B, 3C).

The specialized pancreas-specific functional state of TRM is associated with distinct responses in obesity, shown in our bulk RNA-seq of TRM from pancreas and neighboring tissues. The unique activation of pro-inflammatory and interferon signatures associated with obesity in the pancreatic TRM (Figure 3D, 3E) may reflect their increased coupling with CD11c^+^ macrophages in situ (Figure 2F-2J) and demonstrate pancreas-specific vulnerability to metabolic stress. Higher CD107a^+^GZMB^+^ pancreatic TRMs correlating with BMI link these gene expression changes with enhanced effector functions (Figure 5D). Notably, the changes in pancreatic TRM density and function with rising BMI were not associated with altered TCR diversity (Figure 3G), contrasting with adipose tissue where obesity further restricts TCR diversity^75^.

Pancreatic macrophages likely drive the TRM response to obesity and our results highlight the importance of distinct macrophage subsets in regulating immune homeostasis in the pancreas. The highest surface levels of antigen presentation and co-stimulatory molecules were found on CD11c^high^ macrophages (Figure 1J-1L) enabling them to drive high effector molecule expression in T-cells (Figure 5F, 5G). In contrast the FOLR2^+^ macrophages have high surface PD-L1 and low CD58 (Figure 4G – 4I) which attenuates T-cell activity while their high efferocytosis transcript expression (*MERTK*, *AXL*, *GAS6*) may facilitate dead cell clearance without inflammation^76–78^(Figure 4B). By flow cytometry and imaging, we show how obesity disrupts macrophage homeostasis as regulatory FOLR2^+^CD11c^−^ cells are diminished while pro-inflammatory FOLR2^−^ CD11c^+^ macrophages are expanded (Figure 5A-5C, 6A). Macrophage CD58, which binds to TRM CD2^79^ and drives cytotoxic molecule expression (Figure S5D) may further intensify pancreatic macrophage-TRM interactions with increasing BMI (Figure 1J-1L).

This intensified macrophage-TRM coupling produces metabolically driven inflammatory circuits resistant to resolution. The striking polarization that pancreatic macrophages undergo from monocyte-derived precursors including induction of LAM markers (TREM2, CD9, SPP1) and lysosomal/lipid metabolism pathways with CD11c and T-cell activation molecules (Figure 4B, S4D)^17^, may be driven by lipid stress similar to adipose tissue macrophages in obesity^19,20^. The proliferation of CD11c^+^ macrophages (Figure 4B, 4C, S6E) in pancreas may enable them to displace the more immunoregulatory fetal-derived macrophages (Figure 5A, 5B).

These findings have significant clinical implications. Obesity-related pancreatic diseases may arise via a self-reinforcing cycle of inflammation, damage, and inadequate resolution. Accumulation of CD11c^+^ macrophages and TRM near ductal epithelium (Figure 2B, 2C) suggests possible direct disruption of barrier function and drainage—key pancreatitis initiators^80^. Proliferation of FOLR2^−^CD11c^+^ macrophages and TRM clustering near inflamed Gal-3^+^ epithelium (Figure 6B, 6D) could create cytokine-rich microenvironments that drive KRAS signaling, progression of pre-malignant lesions and genomic instability via chronic exposure to inflammatory mediators (e.g. TNF-α, IL-1β, ROS)^81,82^. These Gal-3-centered niches may link obesity with increased pancreatic cancer risk by promoting pre-malignant transformation^61,83^. Given the recent introduction of pharmacotherapy that reduces adiposity, it will be important for future studies to establish whether weight loss alone can uncouple the pro-inflammatory circuits in the pancreas. Targeting specific cellular interactions within the pro-inflammatory microenvironments (e.g., CD11c^+^ macrophage-TRM coupling, Gal-3-epithelial crosstalk) may help break the chronic inflammation cycles in high-risk individuals.

Collectively, our findings provide novel insights into how obesity reshapes pancreas immune homeostasis through the formation of pro-inflammatory macrophage-T-cell circuits that are centered on inflamed exocrine epithelium. These results provide mechanistic insight into how obesity may drive pancreatic diseases and will inform new strategies to better mitigate chronic pancreatic inflammation.

Despite these advances, our study has limitations. The cross-sectional design limits tracking temporal evolution of inflammatory neighborhoods and determining if immune cell changes are causes or consequences of the inflammation. Longitudinal studies with serial sampling of pancreas tissue are impossible to perform in humans. Although multivariable analysis is useful to adjust for known confounders it cannot account for unknown confounders to definitively establish causality. Also, BMI – although used for diagnosing obesity - does not discriminate between fat and lean body mass or provide information about fat distribution, particularly visceral adiposity which may be particularly important for pancreatic disease pathogenesis.

## ACKNOWLEDGEMENTS

This work was supported by the Louis V. Gerstner, Jr Scholars Program and NIH grant K08 DK122130 to S.W. D.L.F was supported by NIH grant P01 AI106697. A.E. was supported by funding from the German Research Foundation DFG (EI1185/1-1). Research reported here was performed with support of the Molecular Pathology Shared Resource of the Columbia University Herbert Irving Comprehensive Cancer Center (supported by P30 2P30CA013696), Center of Translational Immunology Flow Cytometry Core (supported by grants S10OD030282, S10OD020056, and P30CA01369), the Sulzberger Columbia Genome Center, and the Columbia Single Cell Analysis Core (supported by grant P30CA013696 and P30DK132710) and the Experimental Pathology core facilities, shared resources at the Laura and Isaac Perlmutter Cancer Center (supported by NIH grant CA016087).

The content is solely the responsibility of the authors and does not necessarily represent the official views of the NIH. We wish to thank the donor families for their generosity and the exceptional efforts of the transplant coordinators and staff of LiveOnNY for making this study possible.

---

**Figure S1.**
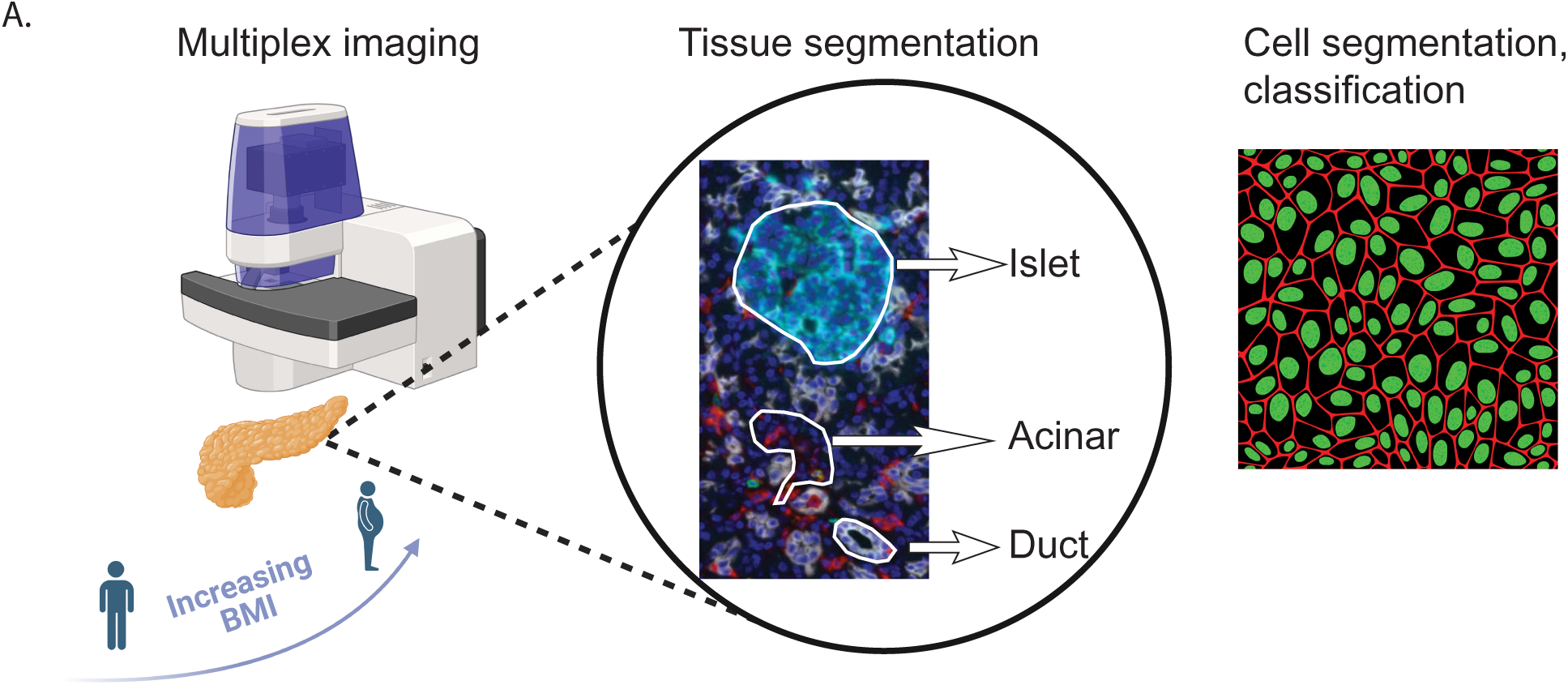
related to Figure 1. Schematic for immune lineage analysis in exocrine and endocrine pancreas of obese and non-obese organ donors. (A) Multiplex imaging was performed on obese and non-obese organ donors, the cytokeratin 19 (CK19) and chromogranin markers were used to segment tissue into ductal, acinar and endocrine compartments, followed by cell segmentation and classification.

**Figure S2.**
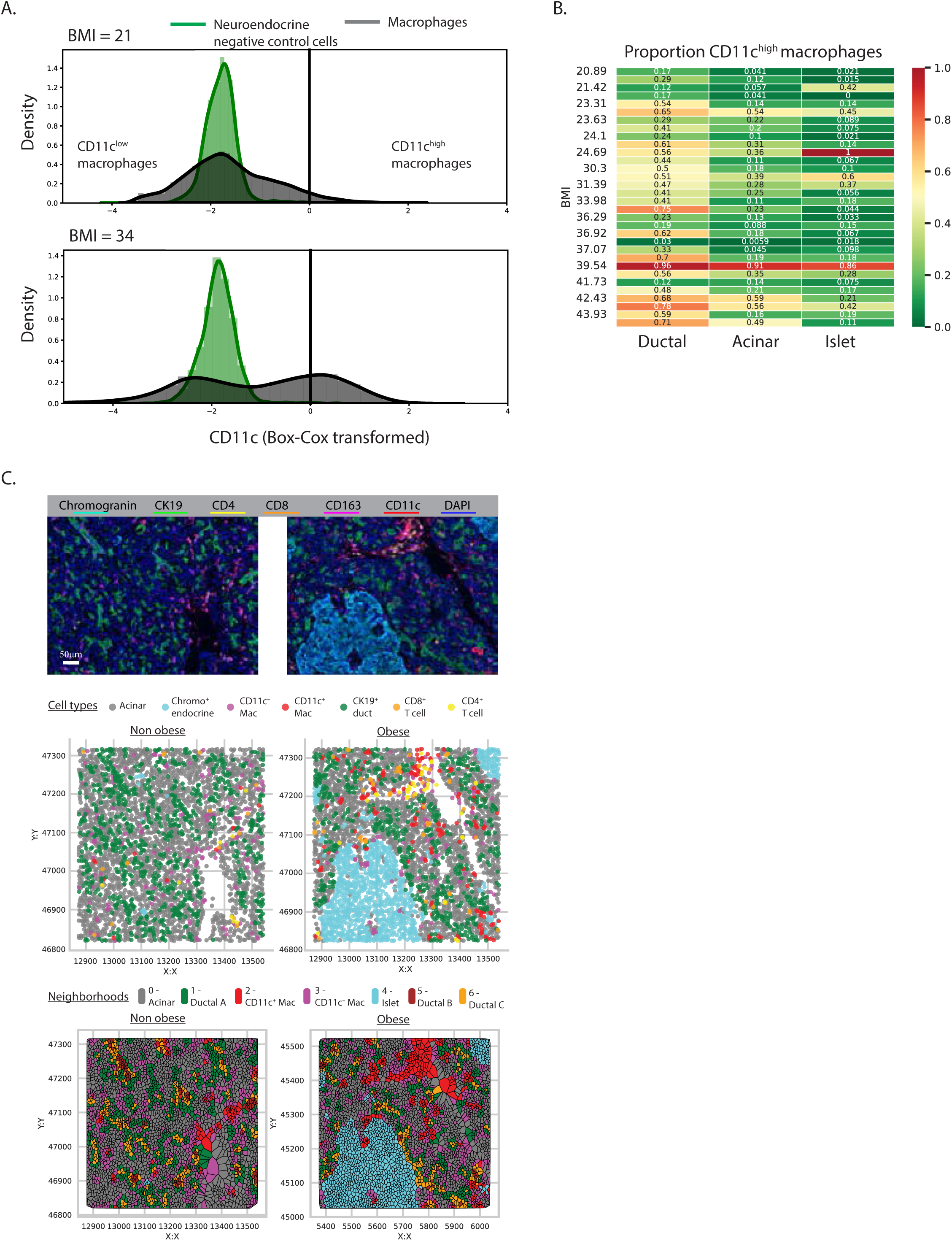
Related to Figure 2. Macrophage classification and neighborhood analysis strategy. (A) Box-Cox transformed density plots illustrating the gating strategy to classify CD11c^high^ and CD11c^low^ macrophage subsets with comparison to CD11c^−^ neuroendocrine cells. (B) Heatmap showing the ratio of CD11c^high^ macrophages to the total macrophage population across pancreatic compartments in donors with BMI between 21 – 47 kg/m^2^. (C) The cell and neighborhood classifications are depicted. Multiplex fluorescence images of pancreatic tissue sections from lean (left) and obese (right) donors, showing CD163^+^ macrophages (red), CK19^+^ ducts (white), chromogranin^+^ islets (cyan), CD4^+^ T-cells (yellow), CD8^+^ T-cells (green), and DAPI-stained nuclei (blue) (top). Based on the multiplex images, the classified cell types (CT) from lean (left) and obese (right) donors are plotted on a Cartesian plane (middle). Voronoi diagrams of these images depicting spatial clustering of cell types into seven neighborhoods (NHs) for lean (left) and obese (right) donors, plotted on a Cartesian plane (bottom).

**Figure S3.**
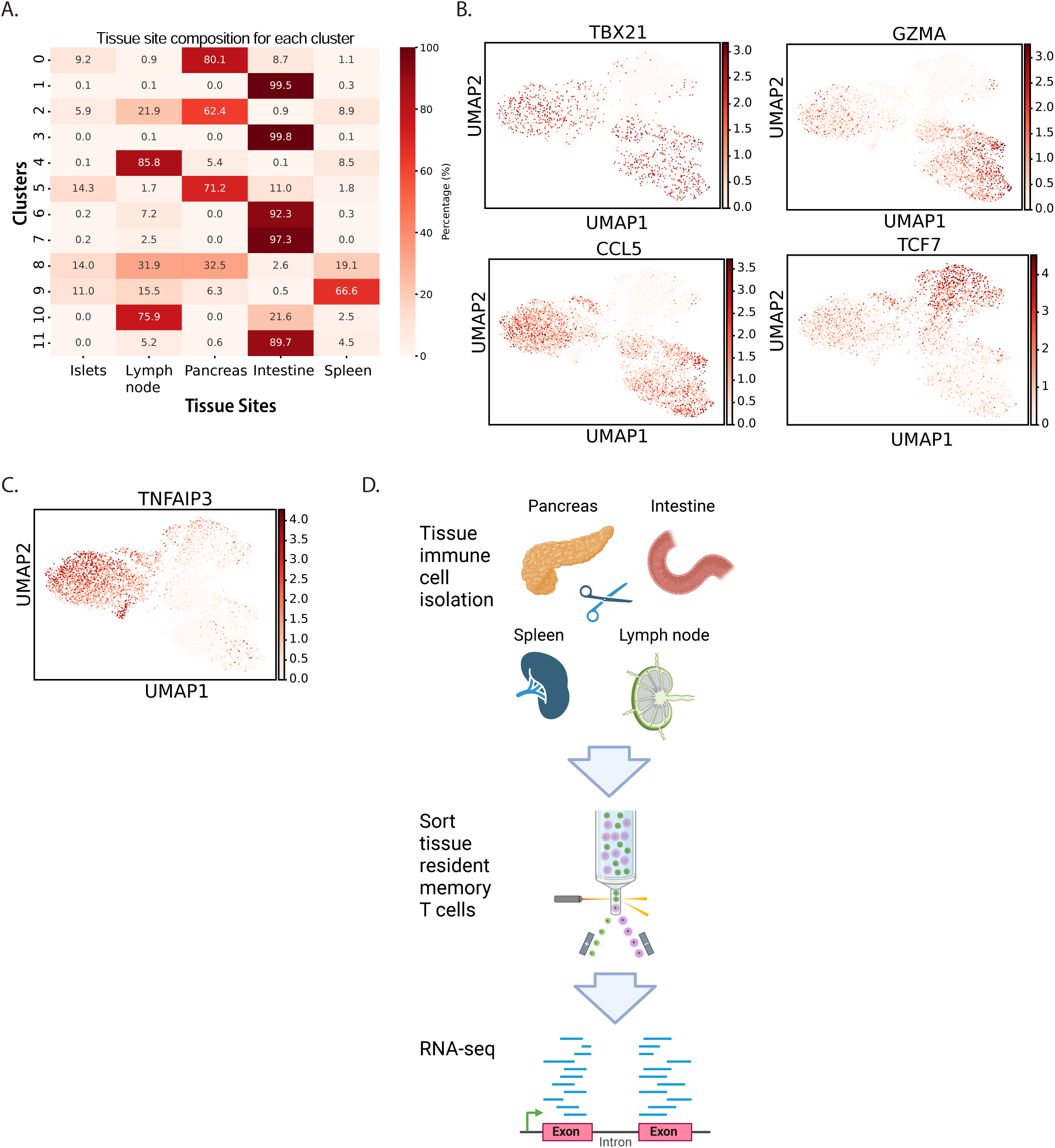
related to Figure 3 Quantification of tissue enrichment for each T-cell cluster in the single cell sequencing dataset, feature plots of selected marker genes for the clusters; and a schematic of the bulk RNA-seq workflow comparing CD8 TRM from obese and non-obese donors. (A) The percentage of cells from each tissue - pancreas, islets, lymph node, intestine and spleen – is shown for each cluster identified from the UMAP embeddings of the CITE-seq dataset. The color scaling is across the rows. (B) Normalized values for the indicated transcripts are depicted on the scaled feature plots including markers of effector memory (*TBX21*, *GZMA*, *CCL5*) and naïve/central memory T-cells (*TCF7*). (C) Feature plot of the negative regulator of inflammation, *TNFAIP3*. (D) Schematic showing the workflow for sorting and bulk RNAseq of CD8 TRM from multiple tissues of obese and non-obese organ donors.

**Figure S4.**
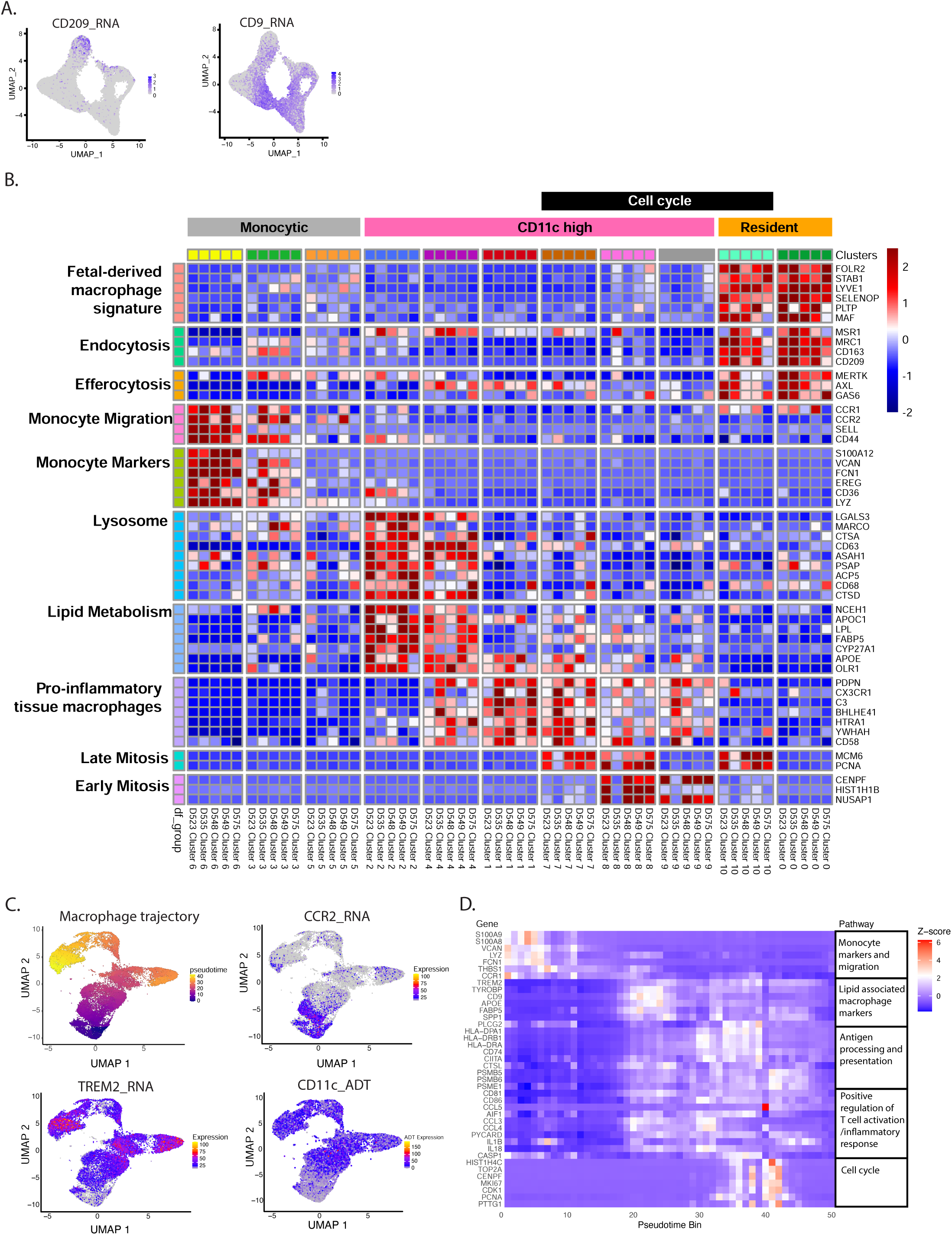
Related to Figure 4. Cluster marker visualization across individual donors and pseudo-time trajectory model of macrophage gene expression. (A) Feature plots show the normalized expression of the *CD209* (left) and *CD9* (right) with expression level corresponding to purple color intensity for individual cells on the UMAP. (B) Shown is a heatmap of normalized RNA expression for marker genes of the macrophage clusters scaled by row and averaged for each individual donor and cluster. The cluster and donor numbers are shown along the bottom with corresponding cluster colors indicated along the top with macrophage classifications based on phenotype and gene expression. The genes are organized by function (left). (C) Series of Uniform Manifold Approximation and Projection (UMAP) plots showing the inferred pseudo-time trajectory of macrophage differentiation (clusters #1-9) and associated expression of key markers that are significantly associated with trajectory progression. The UMAP layout is derived from transcriptional profiles, with cells colored by the following: Pseudo-time progression along the differentiation pathway (top left); *CCR2* RNA (top right); *TREM2* RNA expression (bottom left); CD11c protein expression as measured by antibody-derived tag (bottom right). Color intensity corresponds to the respective feature’s scaled and normalized value. (D) Heatmap of Z-score normalized gene expression (rows) for key marker genes that are significantly associated with pseudo temporal trajectory progression. The pseudo time bins are shown along the bottom and genes organized by function based on gene set enrichment analysis are shown along the right side.

**Figure S5.**
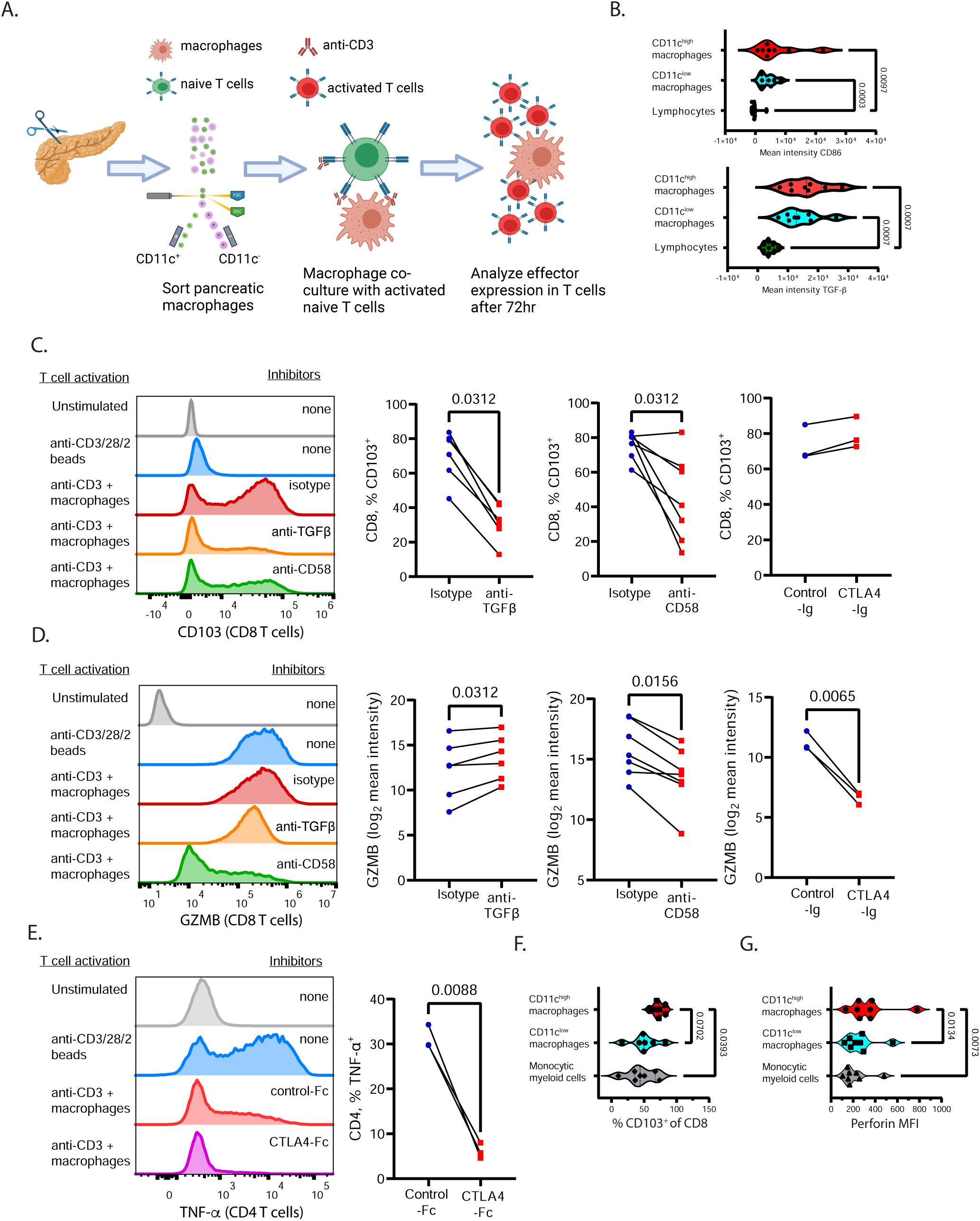
related to Figure 5. The impact of pancreatic macrophage-derived factors on T-cell function. (A) Schematic of the macrophage/T-cell co-culture system used to define how the different pancreatic macrophage subsets influence effector and residency molecule expression by T-cells. (B) Shown are the mean fluorescence values of surface CD86 (top) and intracellular TGF-β (bottom) by flow cytometry from the indicated pancreatic macrophage subsets (n=8). P values were calculated by one-way ANOVA and Holm-Sidak multiple comparison test. (C, D) Representative histograms (left) of CD8 T-cells show CD103 expression (C) and Granzyme B (GZMB) expression (D) under the following conditions: unstimulated (grey); activation with anti-CD3/CD2/CD28 beads alone (blue); activation with monomeric anti-CD3 in pancreatic macrophage co-culture with isotype control (red) and with antibody inhibitors against TGF-β (orange) and CD58 (green). The compiled data (right) show the impact of the indicated inhibitors (TGF-β, n=6; CD58, n=7 and CTLA4-Fc, n=3) on CD8 T-cell expression of CD103 (C) and GZMB (D) after activation and pancreatic macrophage co-culture. (E) Representative histograms (left) of CD4 T-cells show TNF-α expression under the following conditions: unstimulated (grey); activation with anti-CD3/CD2/CD28 beads alone (blue), activation with monomeric anti-CD3 in pancreatic macrophage co-culture with control-Fc (red) and the CD86 inhibitor, CTLA4-Fc (magenta). The compiled data (right) show the effect of CTLA4-Ig compared to control-Fc on CD4 T-cell expression of TNF-α after activation and pancreatic macrophage co-culture (n=3). P values were calculated using a non-parametric paired t-test to determine inhibitor treatment effect versus control. (F, G) Shown are compiled flow cytometry data from naïve T-cells after activation and co-culture with the indicated pancreatic macrophage subsets quantifying the percentages of CD103^+^ CD8 T cells (F) and mean fluorescence intensity of perforin (n=8) (G). P values were calculated by one-way ANOVA and Holm-Sidak multiple comparison test.

**Figure S6.**
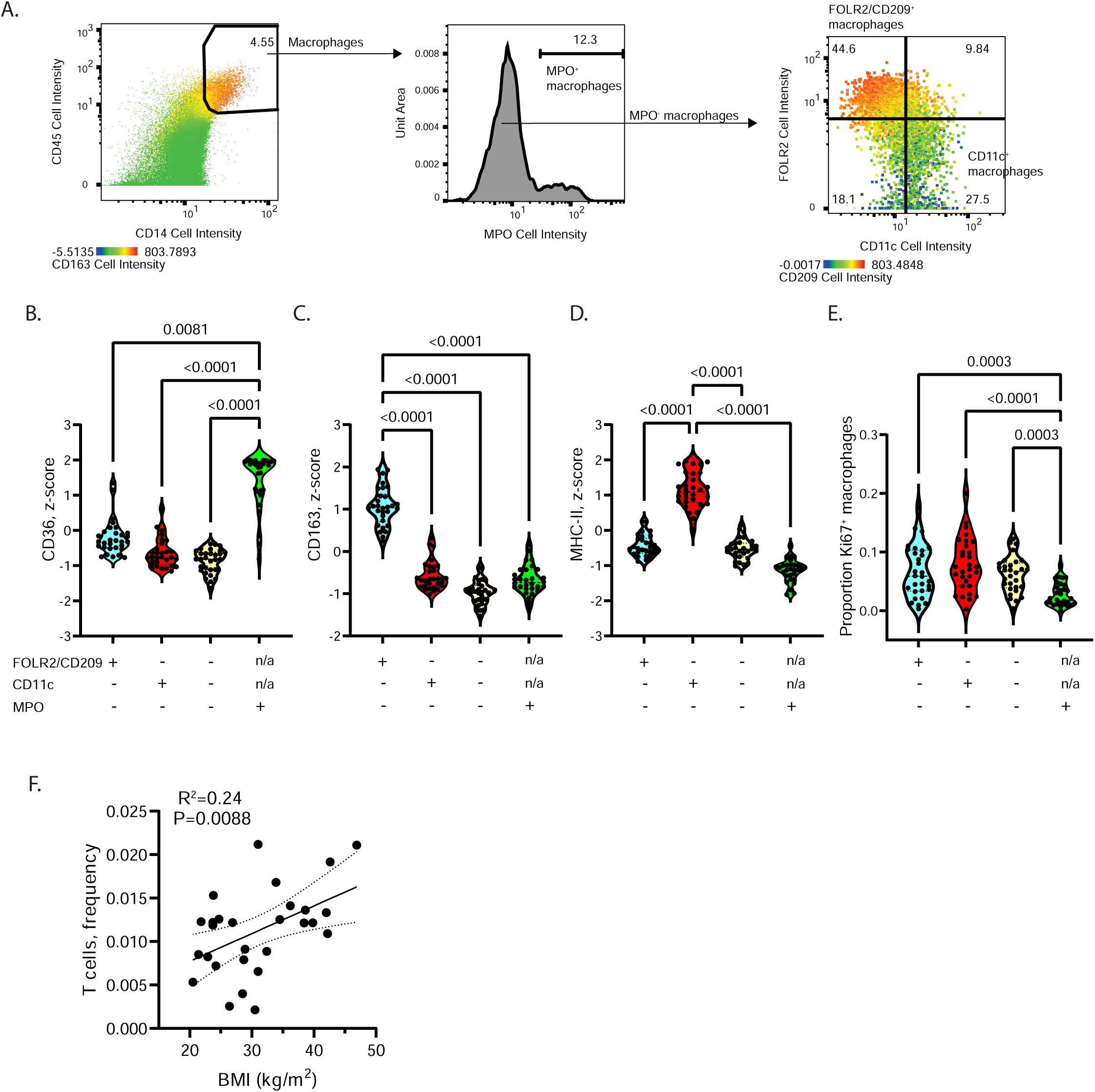
Related to Figure 6. Strategy for macrophage classification based on multiplex imaging of pancreas tissue microarray. (A) Gating strategy for macrophage classification from segmented cells on pancreas tissue microarrays (left), the identification MPO^+^ macrophages (middle), and further sub-setting of MPO^−^ macrophages into FOLR2/CD209^+^ CD11c^−^, FOLR2/CD209^−^ CD11c^+^ and FOLR2/CD209^−^ CD11c^+^ macrophage populations (right). (B-E), Violin plots showing Z score normalized expression of (B) CD36, (C) CD16, (D) MHC-II and (E) the proportion of KI67^+^ macrophages across the indicated phenotypes (shown below each plot). P values were calculated using one-way ANOVA with Dunn’s multiple comparisons testing. (F) The scatterplot of T cells frequency correlated with donor BMI (n=28). The best fit lines, 95% confidence intervals and P values were calculated using simple linear regression analysis.

**Table S1:** Pancreas Disease Risk Factors Examined with Multivariable Models.

|  | Non-obese<br>N=12 | Obese<br>N=20 |
| --- | --- | --- |
| Continuous variables, median (±SD) | | |
| BMI (kg/m<sup>2</sup>) | 23.2 (1.4) | 37.8 (4.7) |
| Age (years) | 49.3 (12.5) | 47.5 (19.3) |
| HbA1c (%) | 5.4 (0.32) | 6.9 (2.5) |
| Categorical variables, n (%) |  |  |
| Diabetes | 0 (0%) | 9 (45%) |
| Heavy alcohol use (2+ drinks/daily) | 1 (8.3%) | 5 (25%) |
| Cigarette use (>20 pack years) ever | 1 (8.3%) | 6 (30%) |
| Male sex | 8 (67%) | 9 (45%) |
| Abbreviations: BMI, body mass index; SD, standard deviation; HbA1c, hemoglobin A1c |  |  |

**Table S2.** Multivariable analysis of pancreas immune cells.

| Variable | CD4 T cells* |  |  | CD8 T cells |  |  | Macrophages |  |  |
| --- | --- | --- | --- | --- | --- | --- | --- | --- | --- |
|  | Regression Coefficient | SE | P value | Regression Coefficient | SE | P value | Regression Coefficient | SE | P value |
| **Acinar** |  |  |  |  |  |  |  |  |  |
| BMI | 0.059 | 0.017 | 0.0012 | 3.63 | 1.28 | 0.0083 |  |  |  |
| Age | n/a | n/a | n/a | 1.26 | 0.66 | 0.066 |  |  |  |
| Diabetes | -0.62 | 0.29 | 0.039 | n/a | n/a | n/a |  |  |  |
| Heavy alcohol use | 0.59 | 0.32 | 0.072 | n/a | n/a | n/a |  |  |  |
| Smoking | n/a | n/a | n/a | 27.53 | 25.54 | 0.29 |  |  |  |
| Male | n/a | n/a | n/a | -35.17 | 22.07 | 0.12 |  |  |  |
| **Ductal** |  |  |  |  |  |  |  |  |  |
| BMI |  |  |  | 3.79 | 1.39 | 0.011 |  |  |  |
| Age |  |  |  | 1.55 | 0.72 | 0.039 |  |  |  |
| Diabetes |  |  |  | n/a | n/a | n/a |  |  |  |
| Heavy alcohol use |  |  |  | n/a | n/a | n/a |  |  |  |
| Smoking |  |  |  | 67.6 | 27.79 | 0.022 |  |  |  |
| Male |  |  |  | -48.84 | 24.02 | 0.0519 |  |  |  |
| **Islet** |  |  |  |  |  |  |  |  |  |
| BMI | 0.73 | 0.27 | 0.010 |  |  |  | 12.67 | 3.85 | 0.0027 |
| Age | n/a | n/a | n/a |  |  |  | n/a | n/a | n/a |
| Diabetes | -6.165 | 4.742 | 0.20 |  |  |  | n/a | n/a | n/a |
| Heavy alcohol use | n/a | n/a | n/a |  |  |  | 126.5 | 76.14 | 0.11 |
| Smoking | n/a | n/a | n/a |  |  |  | 81.64 | 73.65 | 0.28 |
| Male | n/a | n/a | n/a |  |  |  | n/a | n/a | n/a |

Abbreviations: BMI, body mass index; SE, standard error. \*, indicates data are natural log transformed
Results shown are from multiple linear regression models (least squares regression type) in which the outcome variables are CD4 T cell, CD8 T cell and macrophage density within the indicated pancreas tissue compartment. For those analyses that showed significant correlation of BMI with the outcome variable by simple linear regression, multiple linear regression analysis was run with BMI and all the indicated co-variates in an initial model. Only those co variates showing possible association with the outcome variable (P<0.3) were used in the final model. Those variables excluded from the final model due to lack of association with the outcome are indicated (n/a). For each variable the P values reported were calculated using a two-sided test.

**Table S3.** Pathway and ontology analysis of pancreatic TRM transcriptomic signatures.

*Transcripts higher in pancreas-enriched TRM clusters*
| Term | Adjusted P-value | Odds Ratio | Data set |
| --- | --- | --- | --- |
| CTL mediated immune response against target cells | 8.99E-04 | 64.17 | Bioplanet 2019 |
| T-Cell Cytotoxic Mediated Cell Death | 8.34E-06 | 31.53 | Elsevier Pathway Collection |
| Binding of chemokines to chemokine receptors | 3.35E-04 | 22.26 | Bioplanet 2019 |
| T cell receptor regulation of apoptosis | 4.53E-06 | 6.58 | Bioplanet 2019 |

*Transcripts lower in pancreas-enriched TRM clusters*
| Term | Adjusted P-value | Odds Ratio | Data set |
| --- | --- | --- | --- |
| Cellular Response To Corticosteroid Stimulus (GO:0071384) | 1.77E-04 | 33.60 | GO Biological Process 2023 |
| Negative Regulation of MAPK Pathway | 9.66E-07 | 18.01 | Reactome Pathways 2024 |
| Cortisol in Resolving Inflammation | 2.84E-06 | 17.07 | Elsevier Pathway Collection |
| Negative Regulation Of Intracellular Signal Transduction (GO:1902532) | 6.99E-04 | 4.43 | GO Biological Process 2023 |

Abbreviations: TRM, tissue resident memory T-cell
Results shown are from pathway and ontology analysis of differentially expressed transcripts (adjusted P value cutoff < 0.05, absolute value of log<sub>2</sub> fold change cutoff > 0.3) in the pancreas-enriched CD8 TRM clusters (# 0, 2, 5 comprising 3116 cells) compared to intestine-enriched CD8 TRM clusters (# 1, 3, 7 comprising 2686 cells) and all other T-cells in the dataset (6098 cells). Transcripts found to be differentially expressed in the pancreas-enriched CD8 TRM clusters by both comparisons were entered into separate pathway analyses for transcripts that were found to be higher (top) and lower (bottom) in the pancreas-enriched CD8 TRM. For each pathway or ontology term the adjusted P-value, odds ratio and curated data set are shown.

**Table S4.**
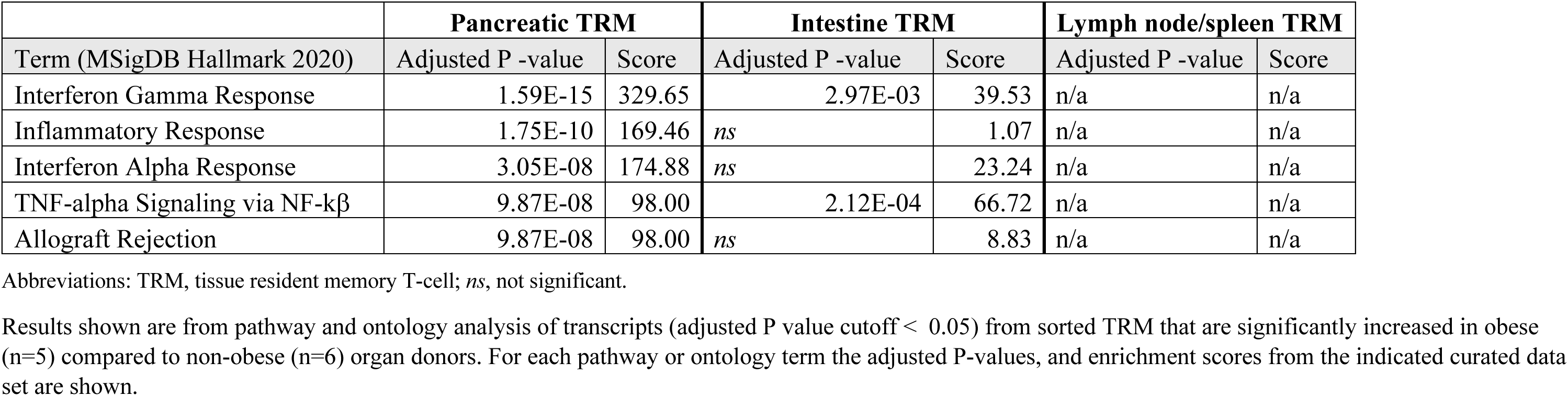
Pathway and ontology analysis of genes from pancreatic TRM that are significantly increased in obesity.

|  | **Pancreatic TRM** |  | **Intestine TRM** |  | **Lymph node/spleen TRM** |  |
| --- | --- | --- | --- | --- | --- | --- |
| Term (MSigDB Hallmark 2020) | Adjusted P -value | Score | Adjusted P -value | Score | Adjusted P -value | Score |
| Interferon Gamma Response | 1.59E-15 | 329.65 | 2.97E-03 | 39.53 | n/a | n/a |
| Inflammatory Response | 1.75E-10 | 169.46 | *ns* | 1.07 | n/a | n/a |
| Interferon Alpha Response | 3.05E-08 | 174.88 | *ns* | 23.24 | n/a | n/a |
| TNF-alpha Signaling via NF-kβ | 9.87E-08 | 98.00 | 2.12E-04 | 66.72 | n/a | n/a |
| Allograft Rejection | 9.87E-08 | 98.00 | *ns* | 8.83 | n/a | n/a |

Abbreviations: TRM, tissue resident memory T-cell; *ns*, not significant.
Results shown are from pathway and ontology analysis of transcripts (adjusted P value cutoff < 0.05) from sorted TRM that are significantly increased in obese (n=5) compared to non-obese (n=6) organ donors. For each pathway or ontology term the adjusted P-values, and enrichment scores from the indicated curated data set are shown.

**Table S5.** Multivariable analysis of pancreatic macrophage subsets.

| Variable | FOLR2/CD209<sup>+</sup> macrophages* |  |  | CD11c<sup>+</sup> macrophages* |  |  |
| --- | --- | --- | --- | --- | --- | --- |
|  | Regression Coefficient | SE | P value | Regression Coefficient | SE | P value |
| BMI | -0.0093 | 0.0037 | 0.020 | 0.0044 | 0.0013 | 0.0028 |
| Age | n/a | n/a | n/a | n/a | n/a | n/a |
| Diabetes | 0.091 | 0.11 | 0.40 | -0.058 | 0.040 | 0.16 |
| Smoking | n/a | n/a | n/a | n/a | n/a | n/a |
| Male | -0.078 | 0.058 | 0.20 | n/a | n/a | n/a |
| TMA1 (ref TMA2) | -0.26 | 0.071 | 0.0012 | 0.16 | 0.023 | 0.0001 |
| TMA3 (ref TMA2) | -0.081 | 0.059 | 0.18 | 0.054 | 0.022 | 0.025 |

Abbreviations: BMI, body mass index; SE, standard error; TMA, tissue micro-array. \*, indicates data are natural log transformed
Results shown are from multiple linear regression models (least squares regression type) in which the outcome variables are the proportions of the indicated macrophage subsets of all macrophages. The FOLR2/CD209<sup>+</sup> subset has the phenotype, CD45<sup>+</sup>CD14<sup>+</sup>MPO<sup>-</sup>CD11c<sup>-</sup> and FOLR2/CD209<sup>+</sup>. The CD11c<sup>+</sup> subset has the phenotype, CD45<sup>+</sup>CD14<sup>+</sup>MPO<sup>-</sup>CD11c<sup>+</sup> and FOLR2/CD209<sup>-</sup>. Multiple linear regression analysis was run using BMI and all the indicated co-variables in an initial model. In addition to the biological co-variables, the analysis corrected for TMA batch. Only those co variables showing possible association with the outcome variable (P≤0.3) were used in the final model. Those variables excluded from the final model due to lack of association with the outcome are indicated (n/a). For each variable the P values reported were calculated using a two-sided test.

**Table S6.**
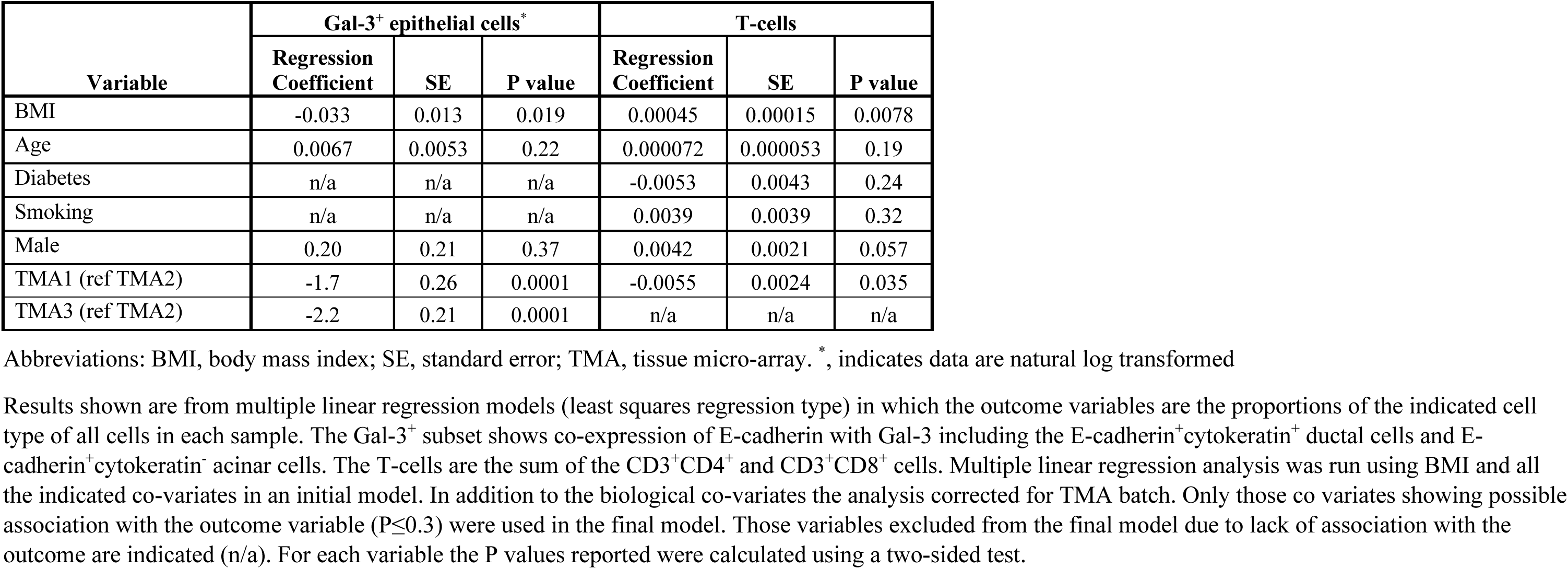
Multivariable analysis of Gal-3+ pancreatic epithelium and T-cells.

| Variable | Gal-3<sup>+</sup> epithelial cells* |  |  | T-cells |  |  |
| --- | --- | --- | --- | --- | --- | --- |
|  | Regression Coefficient | SE | P value | Regression Coefficient | SE | P value |
| BMI | -0.033 | 0.013 | 0.019 | 0.00045 | 0.00015 | 0.0078 |
| Age | 0.0067 | 0.0053 | 0.22 | 0.000072 | 0.000053 | 0.19 |
| Diabetes | n/a | n/a | n/a | -0.0053 | 0.0043 | 0.24 |
| Smoking | n/a | n/a | n/a | 0.0039 | 0.0039 | 0.32 |
| Male | 0.20 | 0.21 | 0.37 | 0.0042 | 0.0021 | 0.057 |
| TMA1 (ref TMA2) | -1.7 | 0.26 | 0.0001 | -0.0055 | 0.0024 | 0.035 |
| TMA3 (ref TMA2) | -2.2 | 0.21 | 0.0001 | n/a | n/a | n/a |

Abbreviations: BMI, body mass index; SE, standard error; TMA, tissue micro-array. \*, indicates data are natural log transformed
Results shown are from multiple linear regression models (least squares regression type) in which the outcome variables are the proportions of the indicated cell type of all cells in each sample. The Gal-3<sup>+</sup> subset shows co-expression of E-cadherin with Gal-3 including the E-cadherin<sup>+</sup>cytokeratin<sup>+</sup> ductal cells and E-cadherin<sup>+</sup>cytokeratin<sup>-</sup> acinar cells. The T-cells are the sum of the CD3<sup>+</sup>CD4<sup>+</sup> and CD3<sup>+</sup>CD8<sup>+</sup> cells. Multiple linear regression analysis was run using BMI and all the indicated co-variates in an initial model. In addition to the biological co-variates the analysis corrected for TMA batch. Only those co variates showing possible association with the outcome variable (P≤0.3) were used in the final model. Those variables excluded from the final model due to lack of association with the outcome are indicated (n/a). For each variable the P values reported were calculated using a two-sided test.

## Material and Methods

### Human Tissue Samples

Tissues were obtained from 68 deceased adult organ donors (Supplemental information 1) as part of organ acquisition for clinical transplantation through an approved protocol and material transfer agreement with organ procurement organizations as described previously^1–6^. All donors had no reported autoimmune diseases, or cancers, tested seronegative for hepatitis B, C, and HIV and did not have a history of chronic pancreatitis or evidence of acute pancreatitis (peak lipase < 3 times upper limit of normal, non-indurated). Use of organ donor tissues does not qualify as ‘‘human subjects’’ research, as confirmed by the Columbia University IRB as tissue samples were obtained from brain-dead (deceased) individuals.

### Isolation and preparation of single cell suspensions from tissue samples

After procurement in the operating room, tissue samples were maintained in cold saline or CoStorSol® (University of Wisconsin (UW) solution (Preservation Solutions, Elkhorn, WI, Cat# PS004) and transported to the laboratory within 2–4 hours. Pancreas and small intestine were carefully inspected for the presence of lymph nodes, and these were removed and processed separately. Spleen, PLN, and small intestine samples were processed using enzymatic and mechanical digestion, resulting in high yields of live leukocytes, as previously described (Carpenter et al., 2018; Gordon et al., 2017; Granot et al., 2017; Kumar et al., 2017; Miron et al., 2018; Senda et al., 2019).

Human pancreatic tissue was processed by the Human Islet Core at Columbia University as described previously^7^. After taking tissue samples for paraffin embedding and image analysis, the body and tail portions of the pancreas were perfused through the pancreatic duct and digested with warmed and circulating enzyme solution in a Ricordi chamber (Biorep Technologies, Inc., Miami Lakes, FL, Cat# RC3–600-MUL) containing Premium Grade Collagenase (SERVA Electrophoresis GmbH, Heidelberg, Germany, Cat#17455.1) and Neutral Protease (SERVA Electrophoresis GmbH, Cat#30301.5) until the majority of the islets were separated from exocrine tissue as examined microscopically by Dithizone-staining (Sigma, Cat#D5130). The pancreas cell suspension, was diluted, washed, and collected using at least 8L of circulating RPMI (Mediatech, Cat# 99–595-CM). Islets were purified using an Optiprep gradient (densities 1.11–1.06) and COBE 2991 blood cell processor, further manually selected to achieve >90% purity by dithizone staining and dispersed using Accutase (Innovative Cell Technologies). Aliquots of the crude pancreas suspension and dispersed islets were immediately cryopreserved in a solution of 90% FBS and 10% dimethyl sulfoxide (DMSO) and stored in a temperature controlled liquid nitrogen freezer for subsequent immune cell recovery.

### Preparation of pancreas tissue for multispectral staining

Representative sections 0.5–1.0 cm in thickness from the pancreas head, body and tail were removed within 12 hours of organ procurement and placed in zinc-buffered formalin (Anatech Ltd.) for 48 hours prior to dehydration and embedding in paraffin by the Columbia University Medical Center Molecular Pathology Shared Resource. For imaging with the 7-color Vectra (Akoya) platform full face sections of the formalin-fixed, paraffin-embedded (FFPE) tissue were used. For imaging with the 32-color PhenoCycler (Akoya) platform, tissue microarrays (TMA) were constructed using 2mm FFPE representative cores (3 cores per donor, and 10 donors per microarray). For all imaging studies 5-µm sections were prepared and immediately processed after drying on positively charged Bond Plus slides (Leica).

### Sorting pancreas immune cells

Cryopreserved tissue suspension aliquots were rapidly thawed at 37°C, washed in RPMI with 10% FBS and DNase I (Sigma, Cat# DN25–5G) (50–100 µg/ml). Dead cells were labeled using Zombie-NIR followed by surface staining with antibodies (Supplemental information 2). For the multi-tissue CITE-seq studies, each different tissue site was labeled with distinct TotalSeq barcoded hashtag antibodies (Biolegend). For sorting, samples were resuspended in PBS with 2% human AB serum and 4’,6-diamidino-2-phenylindole (DAPI) for added sensitivity in detection of dead and dying cells. Sorting was performed using either the BD Influx or FACS Aria Cell Sorters. For multi-tissue CITE-seq, hash tagged cells were sorted from each tissue site at equal proportions. Viable pancreas immune cells were detected as Zombie-NIR^−^ DAPI^−^ CD45^+^. Due to the low abundance of T cells relative to macrophages in pancreas^7^, for the multi-tissue T cell CITE-seq studies, the T cells were further selected as CD2^+^ CD19^−^. Cells were sorted into hanks balanced salt solution (HBSS) with 5% human albumin and 10mM 4’,6-diamidino-2-phenylindole (HEPES) buffer. Post sort viability was confirmed to be >90%. The sorted immune cells were then processed for CITE-seq, bulk RNA-seq and co-culture studies.

### CITE-seq of pancreatic immune cells

After sorting the pancreas immune cells were pelleted, blocked using human TruStain FcX (Biolegend) and then incubated for 30 minutes at 4°C with the TotalSeq-B (10x Genomics Chromium Single Cell 3’ sequencing) or TotalSeq-C (for 10x Genomics Chromium Single Cell 5’ sequencing) Universal cocktail per the manufacturer’s instructions. Immediately after staining and washing 3 times, 8,000–10,000 cells were processed by the Columbia Genome Center Single Cell Analysis Core. Briefly the cells loaded into the 10X Genomics Chromium controller and cDNA synthesis, amplification, and sequencing libraries were prepared using Next GEM Single Cell 5’ or 3’ Kit (10X Genomics) and sequenced on a NovaSeq X (Illumina).

### Bulk RNA-seq of TRM

CD8^+^ TRM (DAPI^−^ CD45^+^ CD3^+^ CD8^+^ CD45RA^−^ CCR7^−^ CD69^+^ cells) from pancreas, small intestine, lymph node and spleen were individually sorted. Analysis was restricted to CD8 TRM because they are the predominant TRM lineage in pancreas. RNA was isolated from cell pellets using the RNeasy Mini Kit (QIAGEN) and quantitated using an Agilent 2100 Bioanalyzer (Agilent Technologies), and library preparation and RNA-seq was performed by the Columbia Genome Center as previously described^4^.

### In-vitro macrophage T-cell co-culture and stimulation

Macrophages (DAPI^−^ CD45^+^ CD14^+^ CD64^+^ CD163^+^ cells) were sorted from pancreas – including the total population and the CD11c^high^ and CD11c^low^ subsets. For TRM co-culture the TRM were sorted from the same sample as the macrophages as DAPI^−^ CD45^+^ CD14^−^ CD64^−^ CD4/8^+^ CD45RA^−^ CCR7^−^ CD69^+^ cells. For naïve T-cell co-culture, peripheral blood naïve T-cells (DAPI^−^ CD45^+^ CD14^−^ CD64^−^ CD4/8^+^ CD45RA^+^ CCR7^+^) were sorted. Macrophages and T-cells were plated together at 1:1 ratio and 50,000 – 100,000 cell density in 96-well U bottom plates in media with 47% RPMI, 47% AIM-V (GIBCO), 6% Human AB serum supplemented with Primocin antibiotic mixture (Invivogen). After resting the cells overnight in co-culture, stimulation was performed using 1 µg/ml monomeric anti-CD3 (OKT3, Biolegend). For analysis of TRM, stimulated cells were incubated at 37°C for 4–6hrs in the presence of GolgiStop (BD Biosciences, cat# 554724) and GolgiPlug (BD Bioscience cat# 555029). For analysis of naïve T-cells, stimulation in co-culture was continued for 72 hours in the presence or absence of inhibitors - anti-CD58 (10 µg/ml, TS2/9, Biolegend), anti-TGFβ (10 µg/ml, ID11, R&D Systems), CTLA-4-Fc (10 µg/ml, R&D Systems), isotype control (10 µg/ml, Mouse IgG_1_, R&D Systems) – followed by 4–6hrs in the presence of GolgiStop and GolgiPlug. Cells were then washed, stained with the fixable viability marker Zombie-NIR, prior to surface and intracellular staining for analytical flow cytometry.

### Analytical flow cytometry

Cells were stained in 5mL polystyrene tubes in the dark using fluorochrome conjugated antibody panels (Supplemental information 2). After staining dead cells using Zombie-NIR, surface staining was done for 30 minutes at 4°C after blocking with human TruStain FcX (Biolegend) in the presence of Brilliant Stain Buffer (BD Biosciences). For intracellular staining, cells were fixed for 60 minutes at 4°C in Cytofix/Cytoperm (BD Biosciences), followed by staining in permeabilization buffer (BD Biosciences Cat) at RT for 30 minutes. Controls were unstained and single-fluorochrome stained cells or UltraComp eBeads (Invitrogen). Flow cytometry data were acquired on an Aurora Spectral flow cytometer (5 laser, Cytek) and analyzed using FlowJo (Treestar, Ashland, OR).

### Multiplex staining on the Vectra platform

The 7-color multiplex panel included DAPI for nuclear counterstaining, and lineage markers for islets, ducts, macrophages and T-cells (Supplemental information 2). Single controls and an unstained slide were stained with each group of slides. After staining, the sections were mounted in Vectashield Hard Set mounting media (Vector Labs) and stored at 4°C for up to 48 hours prior to image acquisition. All samples from 32 donors (Supplemental information 1) were stained together in the same batch.

Multispectral imaging and acquisition at 20x magnification (numerical aperture 0.75) was performed using the integrated Vectra 3 automated quantitative pathology imaging system (Akoya) per the manufacturer’s instructions and as previously described^7^. Representative fields (20-70) from each donor including both exocrine and endocrine regions were captured for analysis.

### Multiplex staining on the PhenoCycler platform

Multiplexed immunofluorescence was performed using a 32-marker panel (Supplemental information 2) on the PhenoCylcer-Fusion platform from Akoya Biosciences, which is based on the CODEX (CO-Detection by InDEXing) method. This technique employs oligonucleotide-tagged antibodies and fluorophores to detect multiple markers through iterative imaging cycles, each followed by an antibody removal step^8–10^. FFPE TMA slides from 28 donors (Supplemental information 1) (3, 2mm cores per donor) were prepared and stained according to the manufacturer’s protocol for PhenoCycler-Fusion (Akoya Biosciences), with the primary antibody incubation extended to an overnight step at 4°C to enhance signal quality ^11,12^. To reduce tissue autofluorescence, a bleaching protocol was applied at the end of the staining process ^13^. Fusion-compatible antibodies were either purchased directly from Akoya Biosciences or generated in-house by conjugating purified antibodies with fluorophore-labeled oligonucleotides using the Antibody Conjugation Kit (Akoya Biosciences, 7000009) (Supplemental information 2).

Imaging was conducted on the PhenoCycler-Fusion imaging system following the manufacturer’s instructions ^14^. The resulting qptiff files were visualized in HALO (Indica Labs). Raw images of the whole slide TMA scans were processed using HALO platform for further analysis.

## QUANTIFICATION AND STATISTICAL ANALYSIS

### Image analysis of multiplex staining using the Vectra platform

#### Tissue and cell segmentation

Images were analyzed using inForm software (inForm 2.3, Akoya Biosciences). Raw 7-plex fluorescence imaging data were pre-processed using the unstained slide and single-color controls for spectral unmixing and autofluorescence mitigation per the manufacturer’s recommendations.

Tissue segmentation was performed as previously described^7^ using inForm software (Version 2.3, PerkinElmer), to classify the pancreas tissue compartments - islets (chromogranin^+^ cell clusters), ductal (CK19^+^) and acinar (highly cellular CK19^−^/chromogranin^−^ areas between the ducts and islets). Cell segmentation was performed based on the DAPI nuclear counterstain and cell classification was performed based on marker combinations as previously described^7^ and as shown (Figure S2). Matrices of individual cells with classifications, X and Y coordinates were exported for further analysis.

#### Macrophage subset classification

We implemented a classification strategy to categorize macrophages into two classes based on CD11c expression levels: macrophages with high CD11c (CD11c^high^ Macs) and low CD11c (CD11c^low^ Macs) expression. Neuroendocrine cells, which lack

CD11c marker expression, served as a negative control. We performed Box-Cox (BCx) normalization on the central signal intensity values of CD11c for both neuroendocrine cells and macrophages to approximate a normal distribution^15^ (Figure S2A) . Histograms of BCx-normalized CD11c intensities revealed that all neuroendocrine cells across all donors consistently exhibited values lower than 0 (Figure S2A). This allowed us to use BCx-normalized CD11c intensity value of zero as a threshold for classifying macrophages as CD11c^high^ (above 0) or CD11c^low^ (below 0).

#### Spatial analysis of macrophage subsets

We quantified the relative abundances of CD11c^high^ and CD11c^low^ macrophages within distinct pancreatic compartments: ductal, acinar, and islet regions. BCx-transformed intensities of CD11c for CD11c^+^ macrophages were calculated and normalized against values from all macrophages (Figure 2C, 2D, 2E). Heatmaps visualized the dynamics of compartmental macrophage composition relative to donor BMI (Figure S2B).

Spatial relationships between macrophages and T cells were analyzed using a custom Python script Single-cell spatial coordinates of each cell were imported into pandas DataFrames for each donor. Pairwise spatial relationships were computed using radius-based nearest neighbor search implemented with scikit-learn’s NearestNeighbors class. A Euclidean distance threshold of 20 μm identified proximal cell neighbors. Tissue architecture was modeled as an undirected graph using NetworkX, with cells as nodes and edges connecting neighboring cells. This approach quantified spatial proximities between T cell subsets (CD4^+^ and CD8^+^) and macrophage subsets (CD11c^high^ and CD11c^low^). The percentages of CD4^+^ and CD8^+^ T cells localized within 20 μm proximity of each macrophage subset was calculated per donor, and Pearson correlation coefficients and p-values were calculated to evaluate relationships with donor BMI.

#### Neighborhood analysis

Neighborhood (NH) analysis was conducted using a modified Python script ^9,16,17^. For each individual cell across all tissue microarrays (4,207,602 cells), was defined a ‘window’ comprising the ten nearest cells based on Euclidean distance. Windows were clustered based on their cell type composition using MiniBatchKMeans (k = 10) implemented in scikit-learn. Seven neighborhood types were identified and validated by overlapping data from fluorescent multiplexed imaging: NH0 (Acinar NH), NH1 (Ductal NH), NH2 (Macrophages CD11c^high^ and T cell NH), NH3 (Macrophages CD11c^low^ and T cell NH), NH4 (Neuroendocrine NH), NH5 (Ductal NH), and NH6 (Ductal and Macrophages NH) (Figure 2I). The frequency of each NH was assessed for each donor, and R-square with corresponding p-values were calculated between donor BMI index and NH frequencies (Figure 2J, 2K).

### Image analysis of multiplex staining using PhenoCycler platform

#### Cell segmentation

Cell segmentation was consistently applied across all samples and was performed based on a three-step process to delineate nuclear, cytoplasmic and membrane compartment using HALO software. First, DAPI was used as the primary nuclear stain to identify DAPI-positive nuclei. A human expert manually annotated representative nuclei by drawing segmentation outlines around them under the classifier tab. The classifier was configured with a resolution of 0.25 µm and a minimum object size of 10 µm², categorizing objects into “background” and “nucleus.” After tuning to confirm segmentation accuracy, the trained classifier was saved for subsequent use. Cytoplasmic regions were extended from the nuclear boundary with a maximum cytoplasm diameter of 1 µm to reduce signal contamination, while membrane segmentation was optimized using dyes displaying a clear honeycomb-like pattern for effective cell border separation, with the software extending the cytoplasm to these boundaries. Single-cell matrices—including average marker expression, with X and Y coordinates were extracted, and all detected cells were exported to FCS files via the “data” tab.

#### Cell quality control classification

Cells were classified using hierarchical gating by a blinded expert using FlowJo (v10.10.0). Cells not expressing sufficient marker signal or not corresponding into defined populations were excluded from cell-type–specific analyses. Data preprocessing and analysis was performed using the workflow described by Windhager et al^18^. The analysis was conducted on an Amazon Web Services (AWS) virtual machine of type r6a.16xlarge (64 vCPUs, 512 GiB of memory, and 25 Gigabit network performance), within a Docker container provided by the Bodenmiller Group (https://github.com/BodenmillerGroup). This containerized setup encapsulated all required dependencies and configurations for the spatial analysis pipeline, enabling seamless and standardized execution.

From a total 3,070,917 segmented cells across 3 tissue microarrays, we delineated 17 cell types based on the combination of the marker expression (Supplemental information 2, 3). For macrophage classification FOLR2 and the CD209 markers were used interchangeably due to inconsistent marker performance between TMA batches. Data from flow cytometry and CITE-seq validated that FOLR2 and CD209 label the same macrophage subset (Figure 4C-4E, S4A). Classified cell types included CD11c^+^ FOLR2/CD209^−^(33,545 cells), CD11c^−^ FOLR2/CD209^+^ (38,453 cells), CD11c^+^ FOLR2/CD209^−^ (16,819 cells), CD11c^+^ FOLR2/CD209^+^ (8,347 cells), CD4 T cells (7,287 cells), CD8 T cells (26,412 cells), Endothelial cells (70,643 cells), Galectin^−^ acinar cells (1,495,446 cells), Galectin^−^ ductal cells (622,864 cells), Galectin^+^ acinar cells (471,434 cells), Galectin^+^ ductal cells (163,449 cells), Islet cells (55,061 cells), Mast cells (13,898 cells), MPO^+^ macrophages (17,992 cells), Neutrophils (17,489 cells), pancreatic stellate cells (PSC) (68,086 cells), and Smooth muscle cells (61,604 cells).

#### Spatial cell proximity graph construction and interaction analysis

To analyze spatial relationships between different cell types, we constructed a custom cell graph in-home Python-based implementation. Single-cell spatial coordinates, corresponding to nuclear centroids, were extracted from the multiplexed immunofluorescence images for each donor. A radius-based nearest neighbor search was implemented using the NearestNeighbors class from scikit-learn to identify proximal cell relationships. For each cell, all neighboring cells within a Euclidean distance threshold of 20 μm were identified as spatial neighbors. The resulting proximity relationships were represented as an undirected graph using NetworkX, where individual cells were represented as nodes, and edges connected pairs of cells within the 20 μm threshold. The graph was constructed separately for each donor to account for donor-specific tissue architecture. Cell neighborhood relationships were formatted into a structured table containing paired cell identifiers and their corresponding donor information and subsequently used for downstream spatial interaction analysis.

Using the spatial graphs constructed above, we assessed spatial interactions between different cell types following the methodology described by Schapiro et al.^19^ . For each pair of cell types, we quantified pairwise interactions to identify significant attractions or avoidances. We calculated an interaction score that represents the degree of spatial association by comparing the observed number of interactions between cells of two types to the expected number under a random distribution hypothesis:

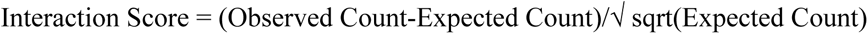

This standardized residual approach provides a normalized measure of interaction strength that accounts for differences in cell type abundances. The resulting scores ranged from approximately -25 (strong avoidance) to +25 (strong attraction), with a score of 0 indicating no spatial preference (Figure 6D). Statistical significance was determined using a permutation-based approach with 1000 iterations, where cell type labels were randomly reassigned while maintaining the spatial graph structure. P-values were calculated by comparing the observed interaction score to the distribution of scores obtained from permutations. Multiple testing correction was performed using the Benjamini-Hochberg procedure to control false discovery rate, with interactions considered significant at adjusted p-value < 0.05. The resulting significant interactions were visualized as a color-coded heatmap, with hierarchical clustering applied to identify patterns of coordinated cellular organization across the tissue. The analysis was performed separately for each donor and subsequently aggregated to identify conserved interaction patterns across the cohort.

#### Local Indicators of Spatial Association (LISA) analysis

Spatial heterogeneity within the pancreatic tissue microenvironment was characterized using Local Indicators of Spatial Association (LISA) clustering, a method adapted from geospatial statistics for single-cell multiplex imaging data analysis and was implemented using the lisaClust package^20^. The analysis was performed on cell type annotations to identify statistically significant local spatial patterns across multiple spatial scales. We applied LISA clustering with k-means parameter k=7 and at three distinct radii (10, 20, 50 µm) to capture both fine-grained and broader spatial relationships For each cell, LISA statistics were computed using a centered local multi-type K-function that measures the proportion of neighboring cells of each phenotype within the specified radii (10, 20, 50 µm), generating a multivariate feature vector representing its local cellular composition. These features were transformed using principal component analysis prior to k-means clustering to reduce dimensionality while preserving variance structure. The LISA algorithm assigned each cell to one of seven spatial context regions (stored as “region” in the data structure), which represented distinct microenvironmental niches within the tissue (Figure 6E, 6F).

### Bulk RNA-seq Analysis

To analyze the effect of obesity on TRM gene expression, RNA-Seq reads from sorted CD8^+^ TRM were mapped, filtered and read counts computed as we previously described^7^. Data from samples that were included in our previous study^7^ (GEO: GSE135582) were merged with 24 additional samples and batch correction was performed using ComBat-seq^21^. For each individual tissue site, principal component analysis and differential gene expression analysis was performed on the adjusted count matrix using DeSeq2^22^ with a design formula to show the effects of obesity on gene expression (Figure 3D, 3E). Gene set enrichment analysis on the differentially expressed genes (DEG) was performed using Enrichr^23^.

### Correlation analysis and data visualization

Simple and multiple linear regression analysis, descriptive statistics of compiled flow cytometry data, graphs and statistical testing were performed using Prism (GraphPad software). Multiple linear regression models were run initially with BMI, age, sex, diabetes, heavy alcohol use (2+ drinks daily), smoking (>20 pack years ever) as independent variables. Those independent variables showing a potential effect on the outcome (P < 0.3) were included in the final model. The plots showing correlation of cells and regions from the three PhenoCycler stained TMAs (Figure 6A, 6C, 6F, S6F) were constructed using data that were corrected for TMA batch based on the multiple linear regression models (Table S5, S6). P values below 0.05 were considered as statistically significant.

### Single-Cell Transcriptomic Analysis of T cells

#### Data Preprocessing and Quality Control

Raw sequencing data were processed with Cell Ranger (v6.0, 10x Genomics)^26^, loaded into Scanpy^27^ (within docker gcfntnu/scanpy:latest) and underwent quality control removing cells with high mitochondrial transcripts (5%) and low unique molecular identifiers (UMIs). Doublets were removed using Scrublet^28^ after score inspection. Post-filtering, 7,550 cells (SW038), 7,788 cells (SW039), and 8,168 cells (SW041) were retained.

#### Hashtag Oligonucleotide (HTO) Demultiplexing

For this multi-tissue CITE-seq study, T cells from different tissue sites were labeled using TotalSeq-C hashtag oligonucleotides (BioLegend). For demultiplexing, raw antibody capture counts were processed separately using Seurat in R. HTO counts were normalized using the centered log-ratio (CLR) method, followed by demultiplexing with the HTODemux function using a threshold at the 99th percentile to remove doublets. Post-demultiplexing, the Seurat object was filtered to include only singlets and were exported as CSV files for downstream analysis.

#### Normalization and Transformation

RNA counts were normalized to 10,000 counts per cell and log-transformed. ADT data were normalized using the centered log-ratio (CLR) transformation. Specifically, ADT counts per cell were log-transformed after pseudo-count addition, and values were scaled relative to the geometric mean expression per cell.

#### Highly Variable Gene Selection and Dimensionality Reduction

Highly variable genes (HVGs) were identified using sc.pp.highly_variable_genes (flavor=’seurat_v3’), testing 2,000–25,000 HVGs. Based on clustering stability (silhouette scores) and variance explained by the top 10 principal components (PCs), 25,000 HVGs were selected. Data were scaled (sc.pp.scale, max_value=10), and PCA (sc.tl.pca) retained 30 PCs.

#### Initial integration and phenotypic filtering

An initial integration was performed on the principal component analysis (PCA) embeddings using Harmony. This step corrected for composite batch effects using a single identifier combining donor (batch) and tissue of origin (tissue_group). The integrated data were clustered using the Leiden algorithm (scanpy.tl.leiden, resolution=1.0). T cells were separated from non-T cell contaminants in the dataset through iterative rounds of clustering and integration. T cell clusters were identified based on expression of canonical markers (CD8A, CD8B, CD2, CD7, IL7R, CD4, CD3E, CD3D) and non T cells were removed based on expression of lineage defining genes (B cells: CD19, MS4A1, CD79A; macrophages: CD14, CD163; mast cells: KIT, TPSAB1) and ADT markers (B cells: Hu.CD19, Hu.CD20.2H7, Hu.CD79b; macrophages: Hu.CD14.M5E2, Hu.CD11b, Hu.CD11c, Hu.CD163; NK cells: Hu.CD56, Hu.CD16).

#### Final integration

To preserve biologically relevant signals from the tissue microenvironment, Harmony was run on the purified T cell dataset to correct for donor-level batch effects only (using batch as the key; max_iter_harmony=20) (Figure 3A). All subsequent analyses were performed on this final, donor-corrected dataset.

#### Visualization of cluster and tissue distributions

Cluster and tissue distributions were quantified using crosstabulation to compute the proportional contribution of each tissue site (Islets, PLN, Pancreas, Small Intestine, Spleen) per cluster in the final donor-corrected dataset normalized by row to percentages (Figure S3A).

#### Differential Expression and Functional Enrichment Analysis

To identify tissue-specific gene signatures, differential expression analysis was performed on the final, integrated dataset. Pancreatic-enriched T-cell clusters (clusters 0, 2, and 5) were compared against all other clusters and against the CD8 predominant small intestine-enriched clusters (clusters 1, 3, and 7) using the Wilcoxon rank-sum test implemented in Scanpy (scanpy.tl.rank_genes_groups, method=’wilcoxon’). P-values were adjusted for multiple comparisons using the Benjamini-Hochberg false discovery rate (FDR) correction. Genes with an FDR-adjusted p-value < 0.05 and a log2 fold-change > 0.25 were considered significantly upregulated. Only DEGs identified in both comparisons are visualized on the heatmap depicting single-cell expression (Figure 3C) and included in gene set enrichment analysis. (Table S3).

### Single-Cell transcriptomic analysis of macrophages

#### Data Preprocessing

Raw sequencing data were processed using Cell Ranger, loaded as Seurat objects and underwent quality control to remove cells with high mitochondrial genes^24^. Each donor sample underwent independent processing with initial dimensionality reduction using buildUMAP and marker identification using findMarkers functions. Multiple rounds of cluster-based sequential selection (2-5 rounds per donor) were performed with comprehensive marker validation including lineage contamination assessment. Clusters were excluded based on high expression of lymphocyte markers (CD2, CD7, CD8A, CD3E, CD19) and mast and epithelial cell markers (SPINK1, KRT8, TPSAB1). Macrophage confirmation utilized established macrophage markers (CD163, CD14, CD33). Multi-modal validation compared RNA-protein expression pairs (CD14/Hu.CD14-M5E2, CD163/Hu.CD163, CD33/Hu.CD33). Final macrophage populations were exported as donor-specific whitelists and saved as cleaned Seurat objects.

#### Data Integration and Batch Correction

Datasets SW031-034 were generated by 3’ sequencing whereas SW037 was generated by 5’ sequencing. The SW031-034 macrophage curated datasets were merged followed by standard processing including NormalizeData, FindVariableFeatures, ScaleData, RunPCA, batch correction using RunHarmony^25^ with “Donor” as the batch variable, then RunUMAP, FindNeighbors, and FindClusters (resolution = 0.7). Initial integration of SW031-034 revealed two minor clusters of non-macrophage contaminants, so these cells were removed. Then the SW037 dataset was added followed by final integration of all 5 datasets using both “Donor” and “Chemistry” as correction variables. The final UMAP was generated using RunUMAP (harmony reduction, dims 1:30), FindNeighbors (harmony reduction, dims 1:30), and FindClusters (resolution = 0.8).

#### Multi-Modal normalization and final selection

RNA data were normalized using log-normalization, while ADT data were normalized using centered log-ratio transformation with normalization.method = “CLR” and margin = 2. Following complete integration, minor clusters 11-14 were excluded due to low expression of macrophage markers, retaining macrophage clusters 0-10 for downstream analysis.

#### Cluster-Specific Differential Expression

Marker genes for each cluster were identified using FindAllMarkers with parameters: only.pos = FALSE, min.pct = 0.25, and logfc.threshold = 0.25 for both RNA and ADT assays using Wilcoxon rank-sum tests. The DEG defining each macrophage cluster were analyzed with Enrichr and curated into 10 functional categories (Figure 4B)

#### Dotplot Visualization

Gene expression dotplots were generated using an analysis pipeline in R with Seurat. Single-cell RNA sequencing data were normalized using LogNormalize with a scale factor of 10,000. The curated gene list of 51 genes organized into 10 functional categories (as described above) was used for visualization. Expression statistics were calculated using a function that computed the percentage of cells expressing each gene (pct.exp) and average expression levels (avg.exp) per cluster. Per-gene scaling was applied across all clusters (0-10), normalizing expression values from 0 to 1 to enable comparison of relative expression patterns. The final visualization combined multiple plot elements using cowplot: (1) a bar chart displaying cell counts per cluster, (2) a dotplot where point size represents the fraction of cells expressing each gene and color intensity represents relative expression levels (navy to darkred gradient), (3) color-coded bars grouping clusters into functional categories. Clusters were ordered as 6, 5, 3, 2, 4, 1, 7, 8, 9, 10, 0 to facilitate biological interpretation (Figure 4B).

#### Donor-cluster matrix generation

For each of the 11 final macrophage clusters (0-10), cells were divided by cluster identity and donor. Average gene expression for each cluster and donor was calculated using AverageExpression with assays = “RNA”. Expression matrices for all clusters were combined to create a final matrix of 71 genes × 55 donor-cluster combinations (5 donors × 11 clusters, Figure S4B).

#### Trajectory Inference with Monocle3

Trajectory analysis was conducted using the monocle3 package^32,33^. The cell data set excluding fetal-derived macrophages (clusters 0 and 10) was preprocessed with principal component analysis (PCA) using 20 dimensions (preprocess_cds, num_dim = 20), followed by UMAP dimensionality reduction (reduce_dimension, reduction_method = “UMAP”). Cells were clustered with cluster_cells, and a trajectory graph was inferred using learn_graph. Pseudotime was calculated with order_cells, with the root cell selected interactively based on biological context. The trajectory was visualized using plot_cells, colored by pseudotime.

#### Differential expression analysis along pseudotime

Differential expression across pseudotime was assessed using graph_test from Monocle3, leveraging the principal graph with 32 cores for parallel computation. Significant genes (q-value < 0.05) were identified, and expression trends were evaluated by calculating slopes via linear regression (lm) using the broom package. Upregulated (slope > 0) and downregulated (slope < 0) genes were filtered per partition, ranked by Moran’s I statistic, and the top genes per category were selected. Expression trends for these genes were plotted along pseudotime with a custom color gradient (blue to yellow, Figure S4C).

#### Pathway Enrichment Analysis and Heatmap Generation

Enrichment analysis^23^ was conducted on the genes significantly correlated with trajectory progression. Representative genes enriched in biologically relevant pathways and their Z-scored normalized values were visualized with a blue-white-red gradient (Figure S4D).

### Software and Environment

#### Computing Infrastructure

Analyses were performed on Amazon Web Services (AWS) virtual machines (r6a.4xlarge with 16 vCPUs, 131 GiB memory, and 12.5 Gigabit network performance; r6a.16xlarge with 64 vCPUs, 512 GiB memory, and 25 Gigabit network performance) and high-performance computing systems with 32 cores. Parallel computation was utilized for computationally intensive analyses including trajectory inference, differential expression testing, and spatial proximity analysis. Containerization was implemented using Docker to ensure reproducibility of computational environments.

#### Image Analysis

Imaging analyses were performed using the PhenoCycler platform (Akoya Biosciences), inForm image analysis software (v2.3, PerkinElmer), HALO platform (v4.0.4, Indica Labs) for multiplexed immunofluorescence processing and single-cell spatial coordinate extraction, and FlowJo (v10.10.0) for cell type classification.

#### Single-cell RNA Sequencing

Single-cell RNA sequencing data were processed using Cell Ranger (v6.0, 10x Genomics^26^), aligning reads to the GRCh38 reference genome. Libraries were prepared using the 10x Genomics Chromium Single Cell 3’ and 5’ Gene Expression platforms (v3 chemistry) with TotalSeq-B and TotalSeq-C antibodies (BioLegend) for CITE-seq analysis and hashtag oligo (HTO) multiplexing.

#### Python Analysis Environment

Python-based analyses (v3.9) utilized the following packages: Jupyter notebook (v6.4.5) for interactive analysis and custom spatial proximity scripts, pandas (v1.3.5) for data manipulation and coordinate processing, scikit-learn (v0.24.2)^34^ including NearestNeighbors class for radius-based spatial analysis, NetworkX (v2.5) for graph-based tissue architecture modeling, Scanpy (v1.9.1)^27^, NumPy (v1.21.0), Matplotlib (v3.5.1), anndata.concat (version ≥ 0.10), AnnData (v0.10) ^29^, and Scrublet (v0.2.1) ^28^ for doublet detection. Python implementations were executed in Docker containers including gcfntnu/scanpy:latest and ghcr.io/yezhengstat/adtnorm:latest for ADT normalization where specified^30^.

#### R Analysis Environment

R-based analyses (v4.0.5-v4.2.0) employed the following packages: Seurat (v4.3.0)^24^, SeuratWrappers (v0.3.0), monocle3 (v3.0)^32,33^ for trajectory inference and pseudotime analysis, harmony^25^ for batch correction, cowplot, patchwork, ggplot2 (v3.3.5), sctransform, dplyr (v1.0.7), Matrix, tidyr, ggplotify, pheatmap (v1.0.12), RColorBrewer, readr, SpatialExperiment, imcRtools ^18^, lisaClust (v1.2.0), mclust, dittoSeq, stringdist, igraph (v1.2.6), broom (v0.7.9) for statistical model tidying, and reshape2 (v1.4.4). R packages were installed via CRAN, Bioconductor (BiocManager), or GitHub (devtools, remotes).

## Data and Code Availability

All analysis code and processed data are available upon request.

## Notes

### Competing Interest Statement

The authors have declared no competing interest.

