## Supplemental information 3 for "Dynamic remodeling of the pancreas immune landscape in obesity"

```
root
└── Whole cells (DAPI^+)
    ├── 1_Neutrophils (MMP-9^+; MPO^+)
    └── Non-neutrophils
        ├── 4_Mast cells (Mast Cell Tryptase^+)
        └── Non-mast cells
            ├── 7_Smooth muscle cells (SMCs) (aSMA^high; Podoplanin^low)
            └── Non-SMCs
                ├── T cells (CD3^+; CD45^+)
                │   ├── 8_CD4 T cells (CD4^+; CD8^-)
                │   └── 9_CD8 T cells (CD8^+; CD4^-)
                └── Non-T cells
                    ├── 10_Islet cells (Chromogranin A^+; Pan-Cytokeratin^-)
                    └── Non-islet cells
                        ├── Macrophages (CD14^+; CD45^+; CD163^+; E-cadherin^-)
                        │   ├── 11_MPO+ macrophages (MPO^+)
                        │   └── MPO^- macrophages
                        │       ├── 12_CD11c^- FOLR2/CD209^+ macrophages
                        │       ├── 13_CD11c^+ FOLR2/CD209^+ macrophages
                        │       ├── 14_CD11c^+ FOLR2/CD209^- macrophages
                        │       └── 15_CD11c^- FOLR2/CD209^- macrophages
                        └── Non-macrophages
                            ├── Ductal cells (E-cadherin^+; Pan-CK^+)
                            │   ├── 17_Galectin+ ductal cells (Galectin-3^+)
                            │   └── 6_Galectin- ductal cells (Galectin-3^-)
                            ├── Acinar cells (E-cadherin^+; Pan-CK^-)
                            │   ├── 16_Galectin+ acinar cells (Galectin-3^+)
                            │   └── 5_Galectin- acinar cells (Galectin-3^-)
                            ├── 2_Endothelial cells (CD31^+)
                            └── 3_Stellate cells (aSMA^+; CD31^-)
```

**Supplemental information 3. Hierarchical classification of pancreas cell types.** Shown is the hierarchical framework used to classify segmented cells from the tissue microarrays that were stained on the PhenoCycler platform using a 32-color multiplex panel.

Supplemental information 3
